# DNA-protein interactions in *Meloidogyne incognita*

**DOI:** 10.1101/2025.07.02.662738

**Authors:** Caroline Bournaud, Alwéna Tollec, Etienne G J Danchin, Yohann Couté, Sebastian Eves-van den Akker

## Abstract

**Background:** The root-knot nematode *Meloidogyne incognita*, is a highly destructive parasite that manipulates host plant processes through effector proteins, affecting agriculture globally. Despite advances in genomic and transcriptomic studies, the regulatory mechanisms controlling effector gene expression, especially at the chromatin level, are still poorly understood. Gene regulation studies in plant-parasitic nematodes (PPN) face several challenges, including the absence of transformation systems and technical barriers in chromatin preparation, particularly for transcription factors (TFs) expressed in secretory gland cells. Conventional methods like Chromatin Immunoprecipitation (ChIP) are limited in PPN due to low chromatin yields, the impermeability of nematode cuticles, and difficulties in producing antibodies for low-abundance TFs. These issues call for alternative approaches, such as dCas9-based CAPTURE (CRISPR Affinity Purification in siTU of Regulatory Elements) that allows studying chromatin interactions by using a catalytically inactive dCas9 protein to target specific genomic loci without relying on antibodies.

**Results:** This study presents an optimized *in vitro* dCas9-based CAPTURE for *M. incognita* that addresses key challenges in chromatin extraction and stability. The protocol focuses on the promoter region of the effector gene *6F06*, a critical gene for parasitism. Several optimizations were made, including improvements in nematode disruption, chromatin extraction, and protein-DNA complex stability. This method successfully isolated chromatin-protein complexes and identified four putative chromatin-associated proteins, including BANF1, linked to chromatin remodelling complexes like SWI/SNF.

**Conclusion:** The optimized *in vitro* dCas9-based CAPTURE protocol offers a new tool for investigating chromatin dynamics and regulatory proteins in non-transformable nematodes. This method expands the scope of effector gene regulation research and provides new insights into parasitism in *M. incognita*. Future research will aim to validate these regulatory proteins and extend the method to other effector loci, potentially guiding the development of novel nematode control strategies.

## Background

### Nematode parasitism and the regulation thereof

Root-knot nematodes (RKN), such as *Meloidogyne incognita*, are among the most destructive plant-parasitic nematodes (PPN), causing significant agricultural losses globally. These nematodes achieve parasitic success by secreting a wide range of effectors - proteins that hijack important host cellular processes (i.e. metabolism, structure, physiology, or immunity), to the benefit of the parasite. These effectors primarily originate from two specialized gland cells (i.e two subventral and one dorsal gland), are ultimately injected into host tissues, and are associated with triggering a complex host-pathogen interaction. Over decades, genomic and transcriptomic analyses have substantially advanced our understanding of nematode effector biology [1, 2, 3, 4, 5, 6]. Numerous developmental stages transcriptomes across evolutionary distinct plant parasitic nematodes indicate concerted and dynamic expression patterns of effector genes throughout the nematode life cycle, suggesting shared and precise regulatory control [1, 7, 8, 9]. Consistent with this global view, a small subset of transcription factors have been found to be strongly co-regulated with effectors over time [10, 11, 12]. Notably, a recent study in the cyst nematode *Heterodera schachtii* established a link between a transcription factor, termed subventral gland regulator (sugr1), and expression activation of effector genes. This transcription factor, along with other highly connected regulators, is expressed in secretory gland cells and appears to be influenced by both nematode biology and plant-derived signals [10]. Despite recent advances, the regulatory mechanisms controlling effector gene expression in PPN remain poorly understood. Developing DNA-protein interaction approaches will be key to uncovering major regulators of parasitism genes and advancing functional genomics in plant-parasitic nematodes.

### Technical hurdles for studying gene regulation in PPN

Progress in deciphering gene regulation in PPN has been hampered by major technical constraints. Unlike model nematodes such as *C. elegans*, *M. incognita* lacks efficient transformation systems, limiting the use of advanced genetic tools like CRISPR for *in vivo* functional studies [13]. Consequently, the community has relied heavily on RNA interference (RNAi)-based research, using either host-delivered or soaking-based approaches to silence nematode genes in a sequence-specific manner [14, 15]. While RNAi technology is pivotal, it can suffer from variable penetrance, stability or specificity [16]. Additionally, transcription factors implicated in effector regulation are often expressed at low levels and/or restricted to specific-cell types, such as gland cells, complicating their study. These difficulties necessitate the development of alternative methods to study chromatin-associated regulatory proteins in PPN.

### Conventional ChIP and its limitations in PPN

Chromatin Immunoprecipitation (ChIP) is a well-established method to study protein-DNA interactions and histone modifications at specific genomic loci and under a physiological context. While ChIP has been successfully applied to *C. elegans* and few other nematodes [14, 17, 18, 19, 20, 21, 22, 23, 24], its use remains limited in PPN. Recently, ChIP based on genome-wide histone immunoprecipitation (IP) has revealed the role of histones modifications for regulation of virulence in *M. incognita*, highlighting the need for precise methods to study the chromatin landscape at the gene locus level, particularly for effector genes [25]. Low chromatin yields, cuticle impermeability and physical structure, as well as the small size and complex tissue organization of *M. incognita,* pose substantial difficulties for isolating high-quality chromatin and in an abundant manner [26, 27, 28, 29, 30]. In addition, it remains challenging to generate high-quality and robust antibodies for such experiments [31]. Conventional ChIP assays require prior knowledge of the target protein, which is often unavailable for many nematode regulators, such as transcription factors. The structural instability of TFs, their low abundance, and restricted subcellular localization further complicate antibody production and limit ChIP applications in PPN. As far as we know, such ChIP methods have not been extended to transcription factors or other regulators in PPN. As a result, more flexible and innovative approaches are needed to investigate chromatin dynamics in these organisms.

### dCas9-based CAPTURE: a related ChIP technique for chromatin regulation studies

The dCas9-based CAPTURE protocol offers a promising alternative to conventional ChIP by bypassing the need for specific antibodies (Fig. 1). It uses a catalytically inactive version of Cas9 (dead Cas9 or dCas9 – unable to cleave DNA), guided by single guide RNAs (sgRNAs), to target specific genomic regions without requiring prior knowledge of regulatory proteins [32]. This flexibility allows for the capture of chromatin-protein complexes at specific loci, which can then be identified through mass spectrometry (MS)-based proteomics (AP-MS) [32]. The method has been successfully applied *in vivo* using conventional cell cultures (e.g cancer cell lines [33]; in *S. cerevisiae* yeast [34]), in *in vitro* systems (e.g 293T cells [35]; S2 cells of *D. melanogaster* [36]) and more recently, in multicellular organisms (e.g. plant species *Betula platyphylla* [37]). It has yet to be tested in PPN. Given the non-transformability of *M. incognita* and the challenges associated with traditional ChIP, *in vitro* dCas9-based CAPTURE provides a viable method for studying chromatin dynamics and regulatory proteins in PPN.

**Figure 1.**
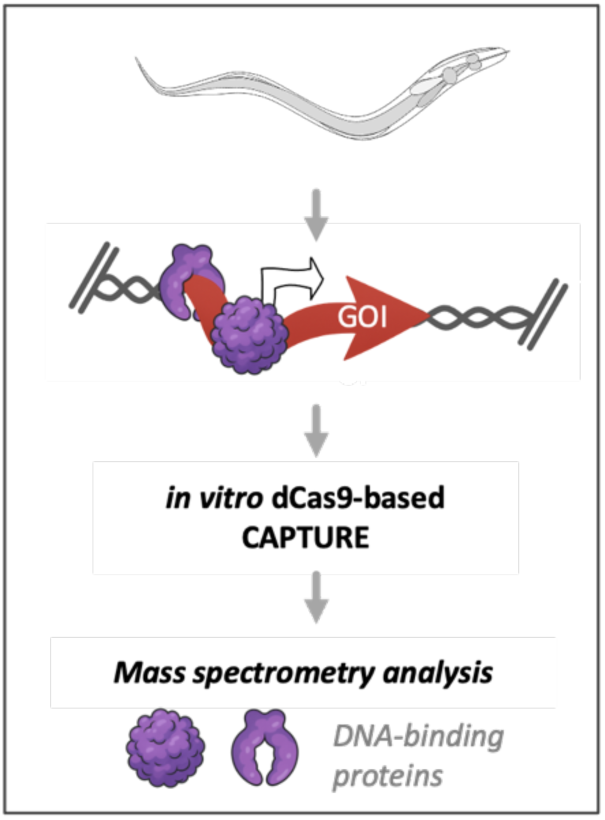
Identification of proteins associated to a specific genomic locus using the *in vitro* dCas9-based CAPTURE technology. The principle relies on a CRISPR-targeted chromatin-based purification strategy, involving the *in vitro* use of biotinylated nuclease-deficient Cas9 (dead Cas9) and synthetic guide RNA (gRNA) incubated with fragmented chromatin from the phytonematode *Meloidogyne incognita*. This recombinant ribonucleoprotein complex, coupled with mass spectrometry (MS)-based proteomics, enables the capture and isolation of proteins associated with the genomic region of interest. GOI : gene of interest.

### Objectives and scope of the study

In this study, we optimized an *in vitro* dCas9-based CAPTURE protocol for *M. incognita*, focusing on the promoter region of the *6F06* effector gene. 6F06 is part of a set of early-expressed effector genes that are transcriptionally coordinated, making it a relevant target for investigating chromatin-based regulatory mechanisms in nematode parasitism [1, 25]. Our work addresses key technical challenges related to nematode physical disruption, chromatin extraction, and purification, ensuring sufficient chromatin yield and stability of protein-DNA complexes. As a proof of concept, we demonstrate the utility of *in vitro* dCas9-based CAPTURE for investigating chromatin dynamics in non-transformable PPN, and pave the way for future studies on regulatory mechanisms in *M. incognita* and other parasitic nematodes of global concern. This approach aims to expand the current understanding of parasitism and contribute to the development of novel strategies for nematode control.

## Results

In this study, we detail major steps necessary for the implementation of the dCas9-based CAPTURE method to identify DNA-associated proteins at a chosen genomic locus of interest in the PPN *M. incognita*. To reach this goal, we made significant improvements to the chromatin preparation protocol, including nematode breakage and homogenization, chromatin fixation and fragmentation. To perform the study of the chromatin-associated proteome of *M. incognita*, we focused on the pre-parasitic juvenile stage, just before the nematode enters the host roots. This stage is optimal for isolating *M. incognita* proteins because it minimizes contamination from plant proteins and allows for easy and abundant collection of biological samples suitable for dCas9-based CAPTURE experiments. For consistency and reliability, we standardized our experiments using a defined number of nematodes, specifically 30,000 pre-parasitic larvae (or eggs, for comparison).

### Evaluation of homogenization techniques

One of key steps in any ChIP assay is isolating a substantial amount of intact chromatin-associated proteome. This process must be adapted in PPN due to their robust cuticle and complex tissue organization. Previous studies on the free-living nematode *C. elegans* have tested various mechanical disruption techniques, including the Dounce homogenizer (two types of pestle with distinct clearance), the Balch homogenizer (which uses a metal chamber and ball bearings with defined clearance) and syringe fixed with a 29-gauge needle (29-G; 0,33 mm x 13 mm), to efficiently break worm tissues and isolate chromatin through repeated passages [25, 29, 30] (Supplementary Fig. S1A). Based on these precedents, we evaluated these different devices as a starting point in optimizing our protocol to disrupt *M. incognita* second-stage juveniles (J2) and recover chromatin-associated proteins. After gentle grinding on ice, microscopic examination indicated that the dounce homogenizer (pestle B) effectively disrupted the *M. incognita* worms, with an optimal disruption time of 7 minutes for complete breakage (Supplementary Fig. S1B). The Balch system proved ineffective for breaking *M. incognita* larvae into pieces, likely due to the nematode’s small size, and was therefore excluded from further optimization steps.

### Isolation of chromatin-rich fractions from nematode lysates

To evaluate the effectiveness of the homogenization devices (dounce homogenizer versus syringe-needle system) in isolating the chromatin proteome from nematode larvae, cell lysates were separated into a soluble fraction (SN) and the crude nuclear extract (P) using biochemical methods (Fig. 2, Supplementary Fig. S2). The soluble fraction (SN) contains cytoplasmic and non-nuclear components, whereas the crude nuclear extract is enriched in nuclear material, including chromatin-associated proteins. Silver-stained SDS-PAGE analysis revealed substantial protein amounts with distinct patterns in the crude nuclear extract (P), with the Dounce homogenizer (pestle B) showing the most pronounced effect. Successful enrichment was confirmed by the visible accumulation of bands likely corresponding to histone proteins in the silver stained gel (P), and immunoblotting for the chromatin marker histone H3. Together, these data suggest that this fraction was enriched in chromatin (Fig. 2, Supplementary Fig. S2). These results confirmed that the Dounce system (pestle B) is effective for both mechanical disruption and recovery of chromatin-associated proteins from worm samples.

**Figure 2.**
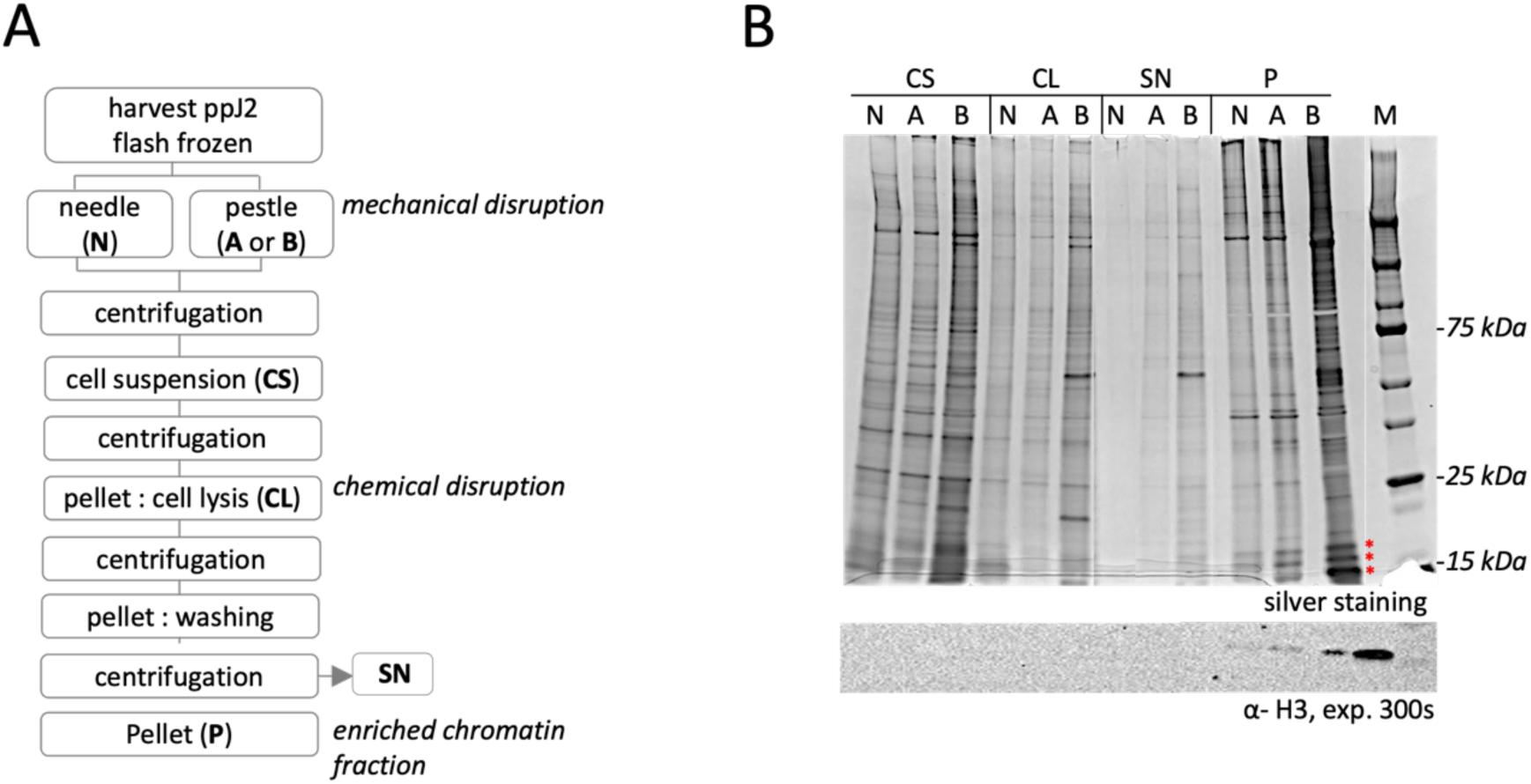
Comparison of homogenisation techniques for chromatin isolation from *M. incognita* second stage juveniles. **(A)** Schematic outlining the steps for chromatin isolation from pre-parasitic juveniles (ppJ2), using two manual homogenization devices: the syringe-needle system (N; 29-G diameter) and the 2 mL Dounce homogenizer (A “type loose“ and B “type tight”). Starting material consists of 30,000 ppJ2 incubated in root extract for 2 hours, followed by flash-freezing in liquid nitrogen and storage at -80°C. Samples from each step were separated into supernatant (**CS, CL, SN**) and pellet (or nuclear crude extract, **P**) fractions. (**B**) SDS-PAGE (4-15%) separation of proteins from the different fractions. Each lane contains protein obtained by the preparation of 500 ppJ2. Protein profiles at key stages of chromatin isolation revealed by silver staining (upper panel). Immunoblotting for histone H3 (bottom panel) highlights chromatin enrichment. Exposure time for chemiluminescence is indicated in seconds (s). Histones presence (≈16 kDA) is marked with red stars, confirmed by the anti-H3 antibody.

### Sonication improves chromatin protein solubilization, but DNA recovery requires further optimisations for ChIP applications

To continue with the chromatin preparation, we applied a sonication step after dounce homogenization to shear chromatin into suitable DNA fragment sizes for downstream ChIP applications. After resuspending the pellet (P), as described in the pipeline (Fig.2), the samples were treated or not by sonication, followed by centrifugation (Fig. 3A). The sonication process favoured the complete disaggregation of the nematode samples into small clumps of cellular debris, as shown in Supplementary Fig. S1B. We then compared the protein profiles of the fractions, with and without sonication (Fig. 3B). Sonication enhanced protein solubilization, particularly for nuclear proteins such as histones, which were visualized in the supernatant fraction (SN, + shear) using silver-stained SDS-PAGE (Fig.3B; upper panel). Notably, histone H3 was detected in the SN fraction only after sonication, confirming the importance of sonication for releasing chromatin-associated proteins into the soluble fraction. However, histone H3 was still predominantly found in the pellet fractions, both with and without sonication step, as confirmed by immunoblotting (Fig.3B; bottom panel). This suggests that the homogenization combined with biochemical treatments followed by the sonication step did not fully release nuclear proteins, leaving some chromatin trapped in the cellular debris. Despite recovering chromatin-associated proteins in this chromatin preparation process, the DNA yield remained insufficient for dCas9-based CAPTURE assays. From 30,000 juvenile nematodes (ppJ2), 100 ng of DNA was recovered, as measured by Qubit, underscoring the limitations of this method for DNA recovery for downstream applications.

**Figure 3.**
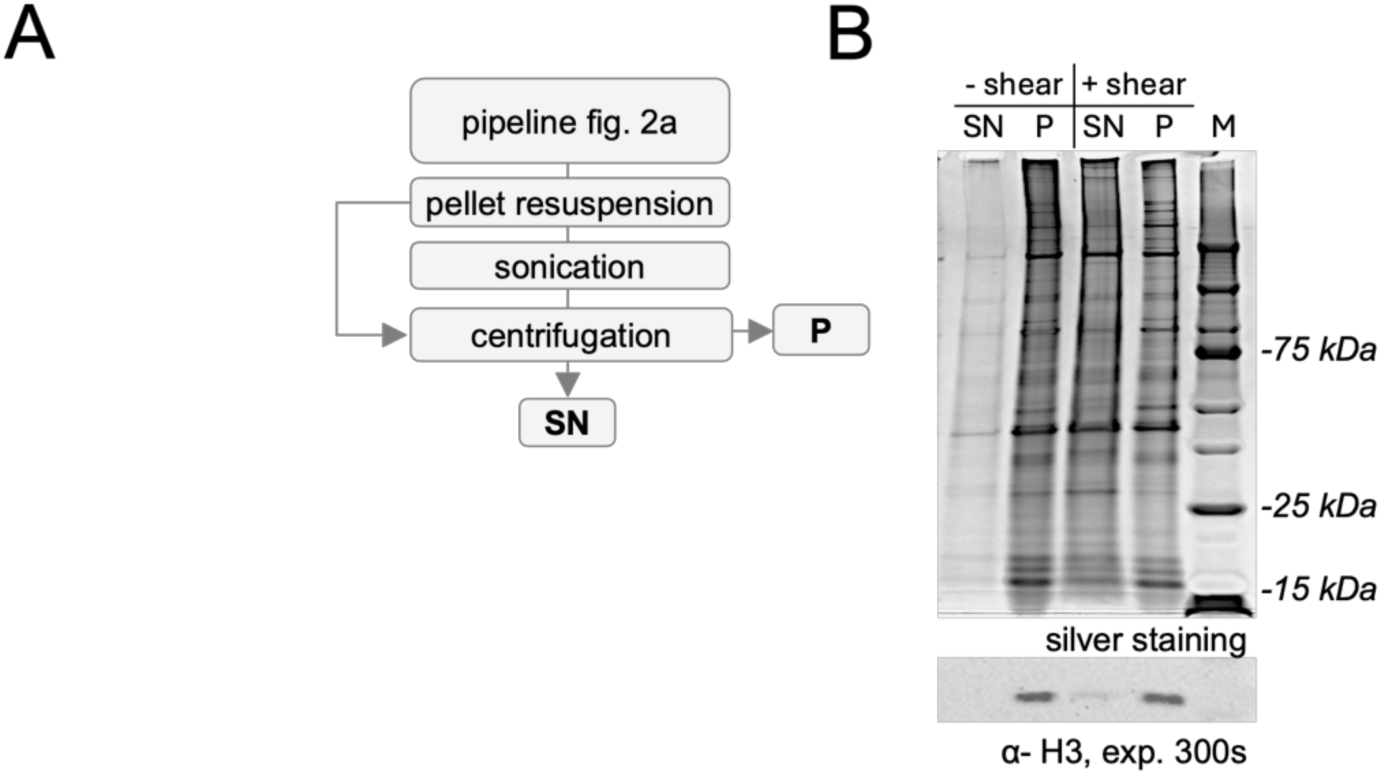
Impact of sonication on chromatin protein solubilization. **(A)** Pipeline for chromatin isolation and shearing following dounce homogenizer with pestle B **(B)** Evaluation of chromatin shearing post-homogenization, where resuspended crude nuclear fractions were subjected to sonication or left untreated. Supernatant (SN) and pellet (P) fractions were collected and analyzed by 4-15% SDS-PAGE (upper panel). Silver staining shows an increase in solubilized proteins in the supernatant after sonication, confirming protein enrichment. Chromatin enrichment was validated by immunoblotting for histone H3 (≈16 kDa) (lower panel). The exposure time for chemiluminescence is indicated in seconds (s).

### Towards improved isolation of chromatin-associated proteins: sonication as a standalone method

Given the low DNA yield and the labour-intensive nature of manual homogenization, we propose an alternative method relying solely on sonication. This streamlined approach offers the potential for both efficient disruption of nematodes and effective chromatin shearing, eliminating the need of manual grinding. A visual inspection (Supplementary Fig. S1B) along with protein electrophoresis and H3 immunoblotting confirmed efficient nematode disruption. Although H3 still remained more abundant in the pellet (P) than in the supernatant (SN) (Fig. 4), the proportion of solubilized H3 in SN was noticeably higher than that obtained with the previous protocol (Fig. 3), suggesting improved recovery of chromatin-associated protein using sonication alone. A similar pattern was observed when eggs were used as the starting biological material (Supplementary Fig. S3). Furthermore, depending on the sonication device, this method allows multiple samples to be processed simultaneously, making it a more time-efficient and scalable option.

**Figure 4.**
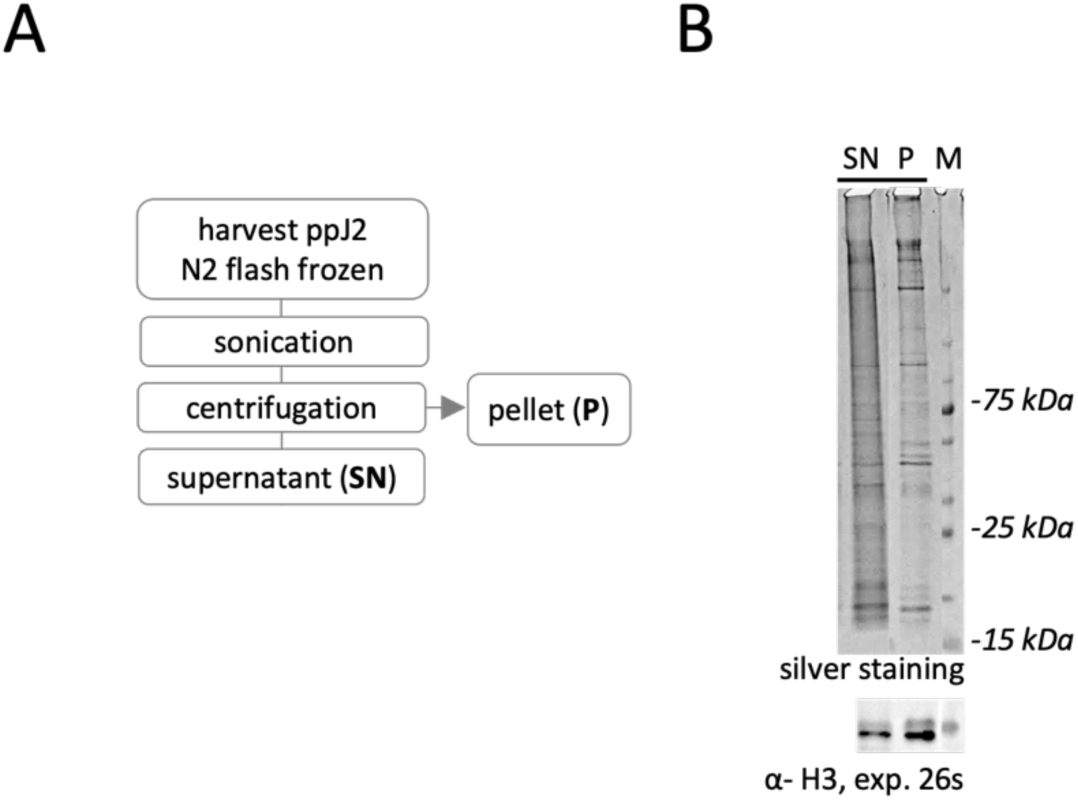
Quick and reliable method to solubilize chromatin-associated proteins of *M. incognita* larvae. **(A)** Pipeline for shearing process and solubilizing chromatin-associated proteins using the Diagenode Bioruptor® as the sonication device. **(B)** Pre-parasitic juveniles were directly sonicated, followed by centrifugation. Aliquots of protein extracts, comprising supernatant (**SN**) and pellet (**P**) fractions, were collected and separated by SDS-PAGE (4-15%), then visualized by silver staining (upper panel). The presence of histones H3 (≈16 kDa) was confirmed by immunoblotting using an anti-H3 antibody to assess the enrichment of chromatin among fractions (lower panel). The exposure time for chemiluminescence is indicated in seconds (s).

### Crosslinking prior sonication for preserving DNA-protein interactions and isolating fragmented DNA in appropriate yield

The chromatin preparation by sonication can potentially disrupt protein-DNA interactions, including those involving transcription factors, highlighting the need for crosslinking prior to shearing. To address this, we conducted an *in-situ* crosslinking step using the conventional fixing agent formaldehyde before sonication. High formaldehyde concentration and/or longer exposure can lead to over-crosslinking, resulting in nonspecific macromolecule associations. To avoid this, we optimized the crosslinking conditions by analyzing the electrophoretic mobility of the proteome using SDS-PAGE. Our results show that crosslinking reactions with either 0,5 or 1% formaldehyde result in similar protein electrophoresis patterns, both distinct from the untreated control (i.e deprived of formaldehyde treatment) (Fig. 5B). The SN fractions, containing solubilized chromatin factors, from both untreated and treated samples were analyzed by DNA electrophoresis to assess fragment DNA lengths and their relative content (Fig. 5C). Although the protein electrophoresis patterns were similar, we found that 0.5% formaldehyde-treated samples resulted in more efficient chromatin shearing, yielding DNA fragments primarily ranging from 100 to 400 bp, nearly identical to the untreated sample. Both untreated and 0.5% formaldehyde-treated samples yielded approximately 1 µg of DNA from 30,000 J2 nematodes, which is twice the amount recovered from the 1% formaldehyde-treated sample. Interestingly, when nematode eggs were used, an apparent well-balanced crosslinking efficiency with minimal protein loss and a suitable DNA size distribution was achieved with 1% formaldehyde for 10 minutes (Supplementary Fig. S4). This suggests that the optimal crosslinking conditions may vary depending on the nematode biological material.

**Figure 5.**
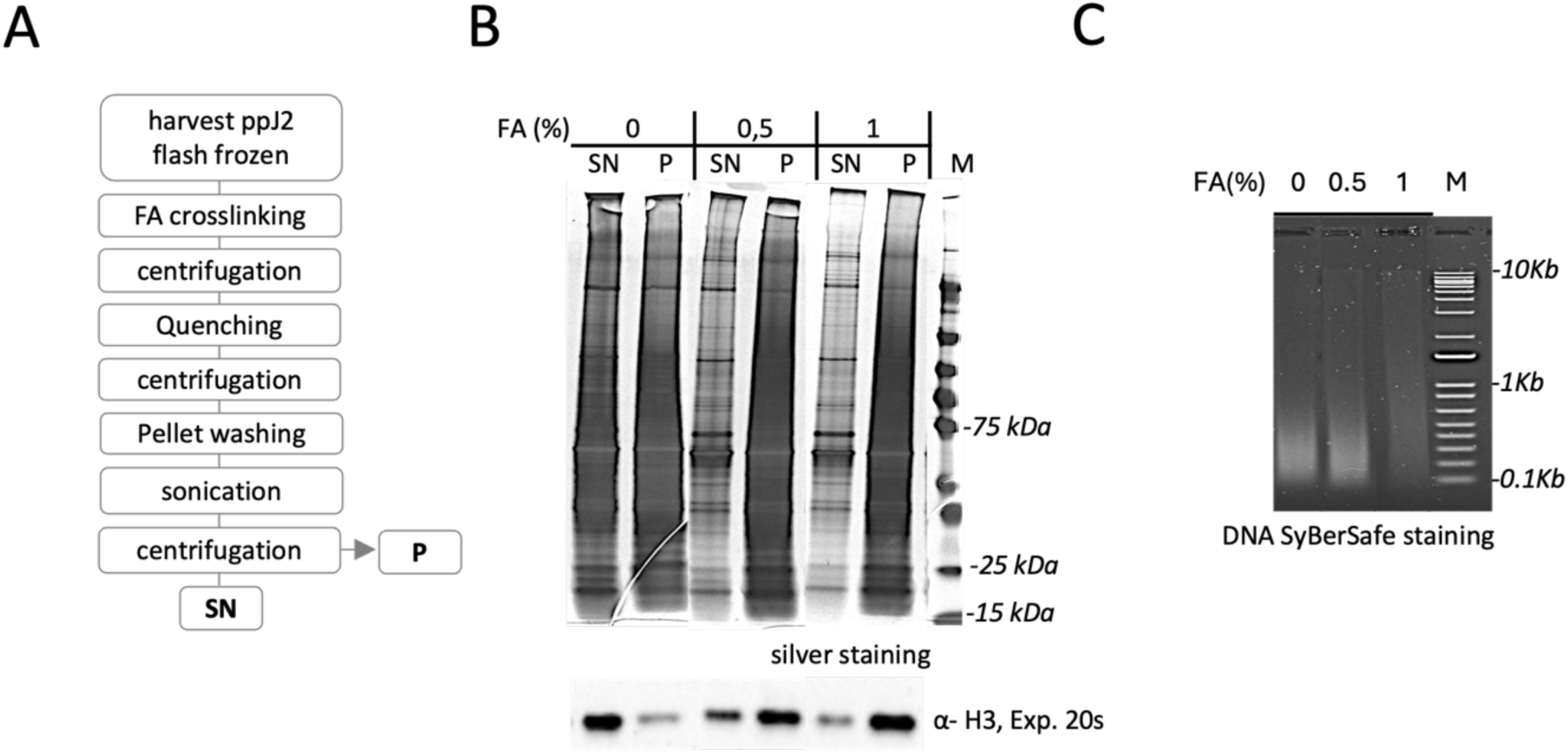
**Influence of crosslinking conditions on chromatin protein solubilization and DNA shearing**. **(A)** Pipeline for chromatin preparation following crosslinking of the larvae, utilizing the Diagenode Bioruptor® as the sonication device. Frozen nematode larvae were crosslinked using formaldehyde (FA), then centrifuged before formaldehyde quenching. Samples were subsequently centrifuged again, and pellets were washed with a physiological buffer. Pre-parasitic juveniles were directly sonicated, followed by centrifugation, and their sheared chromatin profile was analyzed using silver-stained SDS-PAGE and agarose gel electrophoresis. (B) Aliquots of protein extracts, comprising supernatant (SN) and pellet (P) fractions, were collected, separated by SDS-PAGE (4-15%), then visualized by silver staining (upper panel). The presence of histones H3 (≈16 kDa) was confirmed by immunoblotting using an anti-H3 antibody to evaluate the enrichment of chromatin among the fractions (lower panel). The exposure time for chemiluminescence is indicated in seconds (s). **(C)** SN samples generated in (B) were analyzed by SYBRSafe DNA electrophoresis in 2% agarose gel. Fixation at 0,5% FA presents optimal sheared chromatin, where most DNA fragments fall within the size range of 100-400 bp.

### Choice of 6F06 effector as genomic region of interest and design of dCas9/sgRNA in a sequence specific manner

To demonstrate the proof-of-concept for *in vitro* dCas9-based CAPTURE in ppJ2 *M. incognita*, we focused on effector gene promoter regions containing the enriched motif known as the Mel-DOG box [1]. We established criteria for selecting the Mel-DOG genomic region as suitable bait based on previously published results, which indicated: 1) the presence of multiple Mel-DOG boxes within the first 1,000 bp upstream of known dorsal gland (DG) effector genes (considering both position and orientation), 2) Mel-DOG box associated to effector genes exhibiting the highest gene expression levels at pre-parasitic J2 stage; and 3) the ability to design a specific guide RNA (sgRNA) targeting the DNA region of interest. Based on these criteria, we selected the promoter region of the *6F06* effector gene among 34 non redundant effector gene candidates [1]. Consistent with triploidy, three 6F06 homologues are present in the *M. incognita* genome (assembly version V3, 2017 [38]), referring to Minc3s00365g11061, Minc3s08477g42291 and Minc3s02324g29465. Each carries one Mel-DOG motif at - 497 bp or two Mel-DOG boxes from - 504/503 bp and - 347 bp relative to the start codon. The designed sgRNAs were chosen to target specifically a region conserved in the three 6F06 genomic loci, which may improve our enrichment analysis (Supplementary Table S1). The Mel-DOG motifs are positioned approximately 700 bp upstream of the designed sgRNA sequences (located 129 and 139 bp downstream of the start (Met) codon (Fig. 6A). The guide RNAs were designed to bind outside the 6F06 promoter region to avoid steric hindrance between dCas9 and promoter-associated proteins, for which we expected optimal DNA fragments sized around 800-bp (Fig. 6A) to ensure the isolation of related 6F06 promoter proteins of interest.

**Figure 6.**
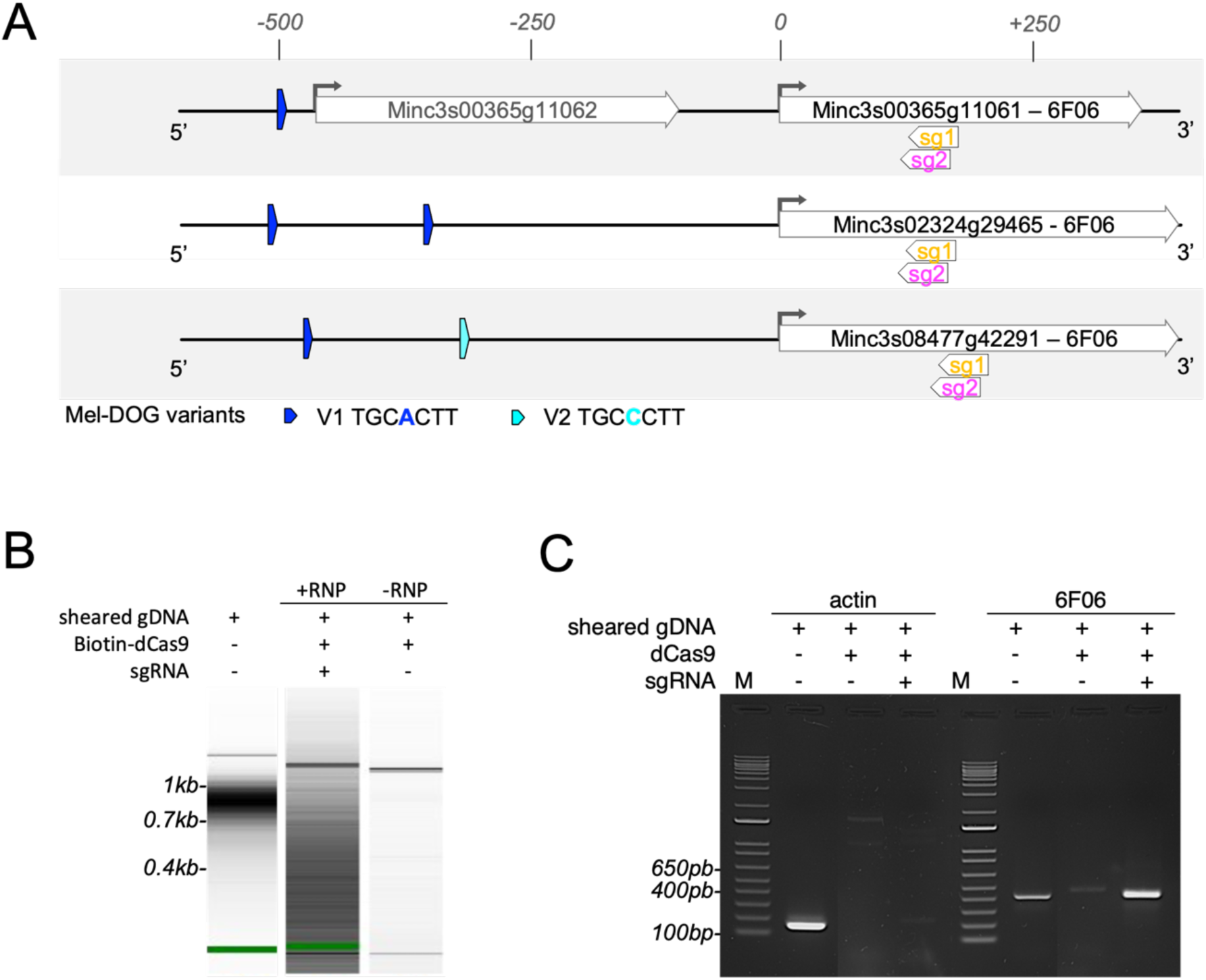
*in vitro* dCas9-based CAPTURE enrichment test using purified genomic DNA of *M. incognita* larvae. **(A)** Map of the 6F06 effector gene loci targeted by *in vitro* dCas9-based CAPTURE. Three genomic regions of the 6F06 gene effector (Minc3s08477g42291, Minc3s02324g29465, Minc3s00365g11061) were used as genomic baits for the *in vitro* R-ChIP assay. The promoters are characterized by the presence of a Mel-DOG box, existing in two variants (blue and turquoise arrows). The 6F06 promoter regions are targeted with two ribonucleoprotein (RNP) complexes corresponding to biotinylated dCas9:sgRNA1 and dCas9:sgRNA2. The selected sgRNAs (sg1 and sg2) are indicated. **(B)** Small fragments of genomic DNA from *M. incognita* larvae were generated by mechanical shearing using the bioruptor®, then incubated with a recombinant dCas9-Biotin loaded with the chosen guide RNAs (sg1 and sg2), or with no sgRNA as a negative control. RNP/gDNA mixtures were added to streptavidin-conjugated resin, followed by washing, and enriched gDNA fragments were pulled down using biotrap affinity purification technology. Enriched DNA fragments were subsequently analyzed using a Bioanalyzer. Gel images showed a broad distribution of DNA fragments targeted by the RNP dCas9:sgRNAs against 6F06 genomic regions compared to the negative control (without sgRNAs). **(C)** PCR analysis of the pulldown DNA fragments from (**B)** was performed targeting the 6F06 promoter regions of interest and Actin as negative control. DNA electrophoresis patterns from PCR demonstrated the presence of 6F06 in the enriched fraction compared to the negative control, while actin was poorly detected in this fraction of interest.

### *In vitro* Cas9/sgRNA cleavage against 6F06 PCR product: validation of the sgRNA choice

To validate the *in vitro* dCas9-based CAPTURE, we first performed an on-target Cas9 cleavage assay to confirm the specificity and efficiency of the sgRNAs targeting the *6F06* promoter region. The *6F06* PCR product (Minc3s02324g29465, 613 bp; Supplementary Fig. S5A; Supplementary Table S2) was successfully cleaved into two fragments using either sgRNA1 or sgRNA2 complexed with Cas9, generating fragments of 241/372 bp and 230/383-bp, respectively (Supplementary Fig. S5B).

DNA fragments isolation in a sequence-specific manner by *in vitro* dCas9/sgRNA system, with the use of genomic *M. incognita* as input – To evaluate the effect of targeting dCas9 to the 6F06 locus, we next conducted the *in vitro* dCas9-based CAPTURE assay using purified genomic DNA. DNA shearing was carried out as described in Fig. 5C, and a bioanalyzer assay revealed an average DNA fragment size of approximately 800 bp (Fig.6B). Fragmented *M. incognita* genomic DNA was incubated with either the biotin-tagged dCas9 / sgRNA complex, and on-target DNA fragments were isolated using a high-affinity Biotrap assay. To assess potential off-target binding by dCas9, a negative control lacking sgRNA was processed in parallel. The dCas9-sgRNA fraction of interest showed significant enrichment and a distinct distribution of dCas9-based CAPTURE DNA products compared to control samples, which included non-targeting dCas9 (without sgRNA) and the input sample. Target DNA regions were analyzed by PCR amplification using primers specific for the 6F06 genomic region. A gene encoding actin was used as a negative control (Fig. 6C; Supplementary Table S2). The *6F06* PCR product was clearly found to be enriched in the + RNP sample compared to the – RNP sample. In contrast, a faint actin signal was detected in the + RNP sample, and was markedly lower than in the input DNA sample, indicating minimal off-target capture. Overall, these results validate the specificity and feasibility of using the *in vitro* dCas9-base capture assay to pulldown a defined genomic locus in *M. incognita*.

### Isolation of chromatin complexes in a sequence-specific manner by *in vitro* dCas9/sgRNA system

We then scaled up the *in vitro* dCas9-based CAPTURE assay using chromatin of *M. incognita* at the ppJ2 stage in a native cellular context (Fig. 7A). Protein and DNA patterns were analyzed from immunoprecipitated fractions treated with dCas9-sgRNA targeting 6F06 genomic loci or non-targeting dCas9. As shown in Fig. 7B, the protein content captured by the *in vitro* dCas9-based CAPTURE was significantly enriched when sgRNAs were used compared to the negative control with no sgRNA. Similarly, chromatin fragments collected from the immunoprecipitated targeting sample exhibited distinct enrichment, with peak sizes around 1,000 bp, compared to the input sample (150-700 bp) (Fig. 7C). To identify 6F06 promoter-associated proteins, purified proteins were analyzed by MS-based proteomics. Among the 193 proteins identified across the analysed samples, 17 proteins were found to be enriched in the +RNP sample compared to the control one (Supplementary Table S3). Among them, five are predicted to localize in the nucleus. To refine our candidate list, we prioritized proteins predicted to be associated with DNA, identifying three putative DNA-binding proteins: a small heat shock protein domain-containing protein (Minc10931a; nuclear, GO:0005634,), a K homology domain-containing protein (Minc3s00047g02560; nuclear, GO:0005634; involved in gene expression regulation, GO:0010468), and the Barrier to autointegration factor (Minc16976; BANF1, DNA-binding, GO:0003677) (Fig. 7D; Supplementary Table S4). These results suggest that the *in vitro* dCas9-based CAPTURE assay successfully identifies potential protein interactants associated with the 6F06 promoter region from *M. incognita* at the ppJ2 stage.

**Figure 7.**
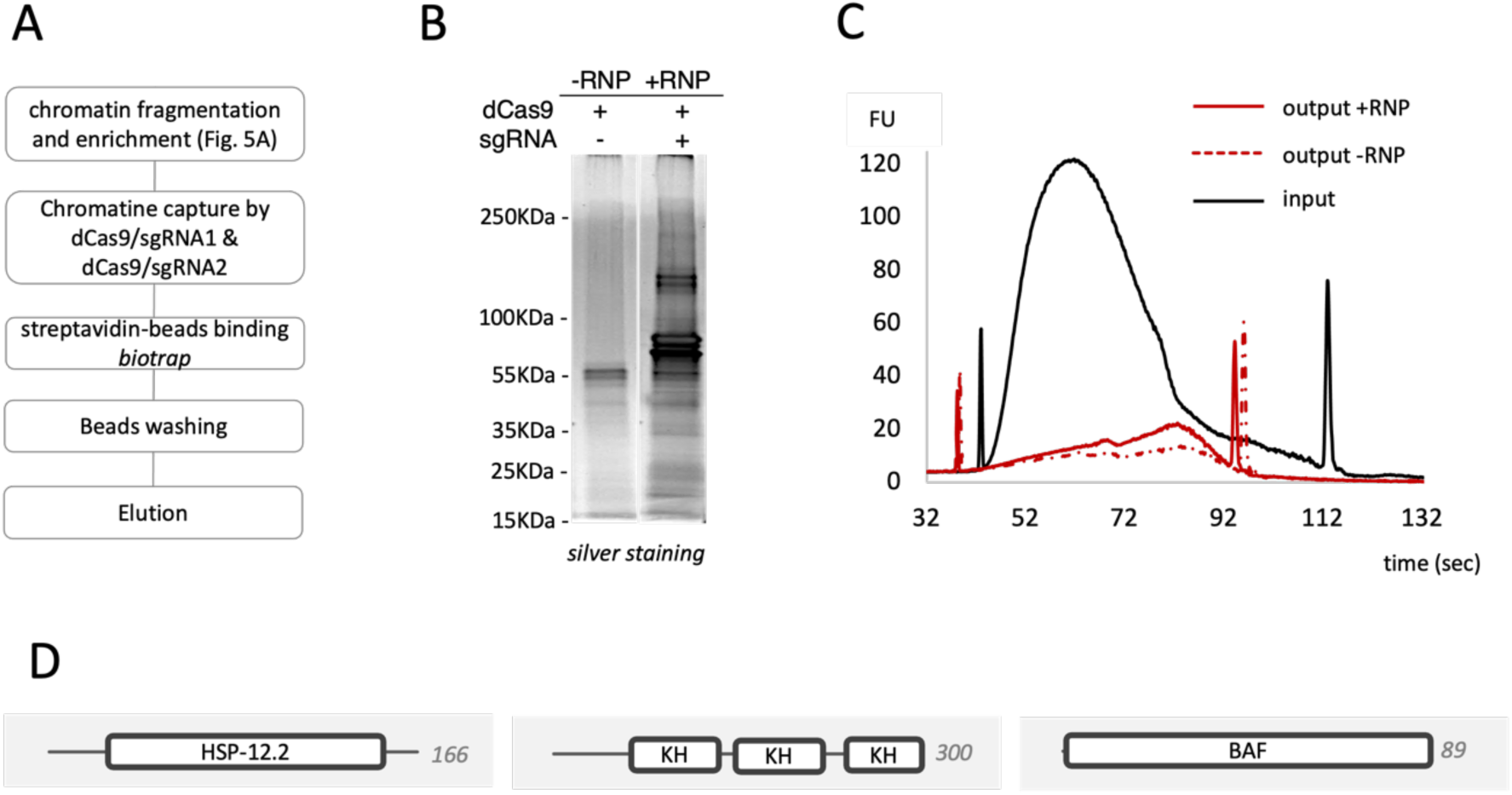
*in vitro* dCas9-based CAPTURE assay under native conditions. **(A)** Overview of the *in vitro* dCas9-based CAPTURE assay using *M. incognita* second-stage juvenile (ppJ2) larvae. Frozen larvae were crosslinked, and small chromatin fragments were obtained by mechanical shearing using the Bioruptor (sonication) of the fixed larvae; The recombinant dCas9-sgRNA complex (+RNP, mixture with equal amounts of sgRNA1 and sgRNA2) were then added to the chromatin mixture, while recombinant dCas9 without sgRNAs (-RNP) served as negative control. Streptavidin-conjugated resin was added to the RNP/chromatin mixture and after washing, enriched chromatin was eluted using Biotrap technology. **(B)** Visualization of elution fractions from the biotrap assay on SDS-PAGE silver-stained gel electrophoresis, showing RNP targeting 6F06 loci (+RNP) and the negative RNP control (-RNP). Several proteins were purified from the eluted fraction of interest. **(C)** DNA size distributions from eluted + RNP and – RNP fractions were analyzed by Bioanalyzer. Sheared and crosslinked chromatin DNA fragments ranged from 200 to 600 bp (Input), while ChIP targeting 6F06 genomic regions or the negative control resulted in fragments from 500 to 800 bp (Outputs). A slight accumulation of DNA fragments was observed in the eluted fraction of interest. **(D)** Interproscan representation of selected proteins identified by mass spectrometry. *In silico* analysis revealed four candidates were potentially involved in chromatin regulation and transcription mechanisms associated with the expression of the 6F06 effector gene during the pre-parasitic stage of *M. incognita*. HSP-12.2 (A0A914L972; SHSP domain-containing protein) BAF (A0A914NCR7; Barrier to autointegration factor; KH (A0A914KPE9; K Homology domain-containing protein). Amino acid length is mentioned on the right side.

## Discussion

### Developing a workflow to study chromatin-associated proteome in PPN - The basis for improvements in order to set up derived ChIP assays

The scope of this study was to evaluate the *in vitro* dCas9-based CAPTURE method for investigating chromatin-associated proteins in *M. incognita*, where genetic transformation remains a challenge. Schematic representation of the method is illustrated in the Figure 8 summarizing several optimizations performed in this study.

**Figure 8.**
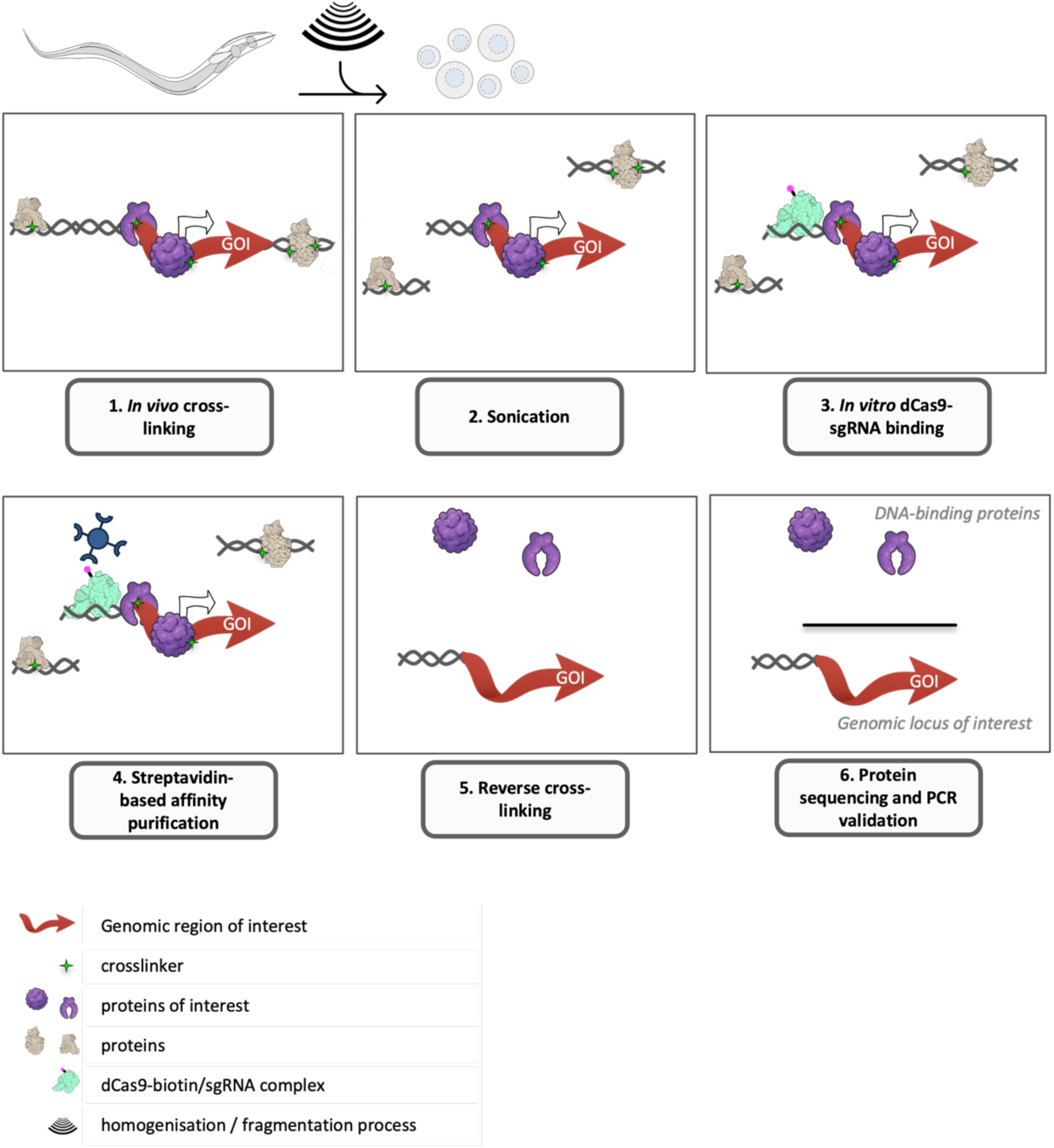
Workflow of the *in vitro* dCas9-based CAPTURE strategy applied to the phytonematode *M. incognita*. Whole nematodes are crosslinked with formaldehyde to preserve protein-DNA interactions. Chromatin preparation involves homogenization and simultaneous chromatin fragmentation using a Bioruptor device. The sheared chromatin is recovered by centrifugation. The dCas9-sgRNA complex is used to target and bind the genomic region of interest. Chromatin-associated proteins are enriched through streptavidin bead capture, followed by reverse crosslinking. Interacting proteins are released, digested with trypsin, and analyzed by mass spectrometry for downstream data processing.

These optimizations included several critical aspects: 1) improving nematode disruption and chromatin extraction, 2) ensuring efficient chromatin fragmentation, 3) employing an appropriate crosslinking condition, 4) refining chromatin capture and 5) improving the signal-to-noise ratio for the *in vitro* dCas9-based CAPTURE assay. We developed a standardized protocol for chromatin isolation from PPN, starting with 30,000 nematode larvae to obtain a yield of 5 µg of fragmented chromatin, making this a feasible and reliable protocol for studying chromatin dynamics and regulation in parasitic nematodes. These advances represent a significant step forward, opening the door to the use of new approaches such as dCas9-based CAPTURE technology for chromatin research in nematodes.

### Challenges in studying chromatin in nematodes - Adapt nematode disruption and chromatin isolation to PPN

Previous studies in the well-established model system *C. elegans* have provided key insights in optimization of chromatin preparation, but adapting these protocols to *M. incognita* comes with unique challenges [18, 28, 29, 39]. Unlike conventional cell cultures, nematode cells are highly heterogeneous, making homogenization and collection of nuclear extracts from all cell types more challenging. Additionally, *C. elegans* and other nematodes generally have much lower chromatin content per unit of body weight, sometimes up to 100 times less than that of mammalian cells [28]. This low chromatin yield makes experiments like dCas9-based CAPTURE assays significantly more difficult. Moreover, *M. incognita*, particularly in its pre-parasitic J2 (ppJ2) stage, has a highly impermeable and resistant cuticle, which hinders the penetration of chemical like fixation agents and complicates mechanical grinding [26, 27]. While manual homogenization using the dounce homogenizer effectively disrupted nematodes’ bodies, it produces low yields of chromatin, making the process labor-intensive and less efficient. Compared to *C. elegans* (typically studied in gravid adults), *M. incognita* ppJ2 are roughly half the size. The size difference, coupled with the nematode’s cuticle properties, necessitated specific adaptations in chromatin extraction procedures. Such modifications were critical to improve chromatin recovery and quality for downstream applications like dCas9-based CAPTURE assays.

### Recovery and integrity of chromatin-associated proteome using different standard devices

Cellular homogenization is a critical step in dCas9-based CAPTURE assays, which involves the disruption of various cellular components (as well cell-, nuclei- or chromatin levels). One of the main challenges in dCas9-based CAPTURE assays is ensuring effective homogenization - disrupting the nematode cuticle while preserving chromatin integrity. We evaluated various homogenization methods, including Balch and Dounce homogenizers, as well as needle/syringe systems. However, these manual techniques led to significant nematode material loss and low chromatin recovery. While Balch homogenization has been successfully applied in *C. elegans*, it proved less compatible with the smaller nematode *M. incognita* [29, 30]. Manual dissociation, although feasible, required large quantities of nematodes due to sample loss, which impacted reproducibility, especially with low-input material. Additionally, higher densities of nematodes increased the mechanical effort needed, making the process time-consuming and raising the risk of destabilizing chromatin-protein complexes. We thus turned to sonication as an alternative, where the chromatin fragmentation step occurs concurrently with homogenization. Sonication conditions were optimized to both break down nematode tissues and fragment chromatin to sizes compatible with downstream dCas9-based CAPTURE assays. Several parameters were critical to efficient and uniform sonication, including preventing nematode sedimentation, which was more pronounced during the early stages of sonication when tissue disruption was minimal. Further optimal conditions of chromatin preparation may be necessary, such as adjusting fine-grained cycle intervals to ensure resuspension of the nematodes between rounds of sonication, and the sonication time and number of cycles or the dounce homogenization protocol to improve chromatin release. Combining sonication with benzonase nuclease digestion proved beneficial for minimizing the destabilization of protein complexes [40].

### Crosslinking considerations to preserve chromatin integrity

In our experiments, we used histone H3 as a chromatin marker due its high conservation across species, allowing us to use commercially available antibodies. Despite stringent sonication conditions, nucleosome interactions with genomic DNA were robust enough to withstand disruption [41]. We also considered immunodetection of DNA polymerase II (PolII) to assess chromatin complex stability [42]. However, the antibodies tested failed to produce consistent results. To further preserve chromatin integrity, we employed crosslinking prior to sonication, a step that required optimization based on tissue type, as different responses to the crosslinking agent (FA, formaldehyde) were observed between egg and juvenile nematodes. Western blot analysis revealed decreasing amounts of histone H3 in chromatin fractions following formaldehyde treatment, suggesting incomplete lysis or shearing, combined with the crosslinking effect. Particular care must be taken to avoid artefactual crosslinking and to ensure the quality of chromatin fragmentation. Importantly and as previously reported, our data confirm that the addition of cross-linkers improves stability without prohibiting dCas9:sgRNA to anchor on chromatin. Thus, both sonication and formaldehyde crosslinking conditions require careful optimization for effective chromatin preparation regarding the biological sample tested [42, 43].

dCas9-based CAPTURE, like other ChIP-derived methods, offers the advantage of isolating chromatin-protein complexes under native physiological conditions. However, enrichment may be insufficient or nonspecific. Although the PCR test targeting the actin gene is indicative, other gDNA loci may have been co-purified. To improve specificity, output DNA sequencing could be included in sgRNA screening to select the most specific guide RNAs. Importantly, we cannot control from which cells the DNA is isolated, and so a mixture of chromatin states is likely. Coupled with the low abundance of DNA-binding proteins (DBPs), particularly in gland cells, this may limit the discovery of parasitism-related gene expression regulators. Factors such as overly stringent chromatin extraction, insufficient crosslinking, and limited protein sequencing depth may have impacted protein enrichment. Nonetheless, dCas9-based CAPTURE provides flexibility and specificity, enabling the capture of chromatin-protein complexes at single or multiple loci, depending on the sgRNA design.

### Proof of concept of dCas9-based CAPTURE: case study to identify proteins putatively associated to 6F06 effector loci

A significant challenge remains in identifying parasitism gene expression regulators in *M. incognita*. The scarcity of secretory gland cells - the primary but not exclusive source of effector proteins - makes it difficult to isolate sufficient quantities for comprehensive identification of chromatin-associated proteomes involved in parasitism. These cells include only one dorsal gland cell and two subventral gland cells, among the approximately 1,000 cells in the nematode (using *C. elegans* as reference, which contains 959 somatic cells). Beyond its technical aspect, dCas9-based CAPTURE tool was expected to provide a comprehensive overview of protein candidates that help elucidate how parasitism regulation occurs in the root-knot nematode *M. incognita*. In our study, the promoter region of the 6F06 effector gene was selected as a DNA bait to conduct dCas9-based CAPTURE experiments coupled to MS-based protein analyses, which identified candidates carrying a DNA-binding protein domain. However, these promising candidates associated with promoter regions of interest, remain to be experimentally characterized.

From our dCas9-based CAPTURE experiments, we shortlisted three candidate proteins targeting 6F06 genomic loci. These included Barrier-to-autointegration (BANF1, BAF1) and two other proteins related to transcriptional regulation, a small heat shock protein (SHSP) domain-containing protein and K homology (KH) domain-containing protein. Although limited data are available on the role of BANF1 in parasitism, it has been shown to interact with SWI/SNF complexes [44]. BANF-1 can act as an epigenetic mediator, either activating or repressing gene expression [45]. Although we did not experimentally verify the role of these proteins in parasitism, several studies suggest a potential role for SWI/SNF complexes in the regulation of parasitism genes, as demonstrated in *Plasmodium falcifarum,* the malaria parasite [46]. A recent review summarized how SWI/SNF complexes are recruited in fungal growth and stress responses, requiring the interplay of multiple transcription factors and covalent histone modifiers [47]. These findings mirror with the previous work in *M. incognita,* which highlighted that approximately 20% of effector genes are regulated by histone modification dynamics. For instance, 6F06 is one of them that exhibits repressive expression during the transition from eggs to juvenile stages (J2), and this repression is associated with H3K9me3 mark [25]. Overall, it would be valuable to further study their functional roles to evaluate their significance in parasitism at the early stages of infection.

The dCas9-based CAPTURE technology has proven to be a useful tool in several biological contexts, offering a deeper understanding of chromatin interactions. The dCas9-based CAPTURE system not only captures chromatin but can also isolate locus-specific long-range DNAs or chromatin-associated RNAs, such as long non-coding RNAs (lncRNAs), which are known to act as trans-regulatory elements in concert with regulatory proteins to control gene expression [48]. Evidence from the literature shows that SWI/SNF complexes require guidance from transcriptional regulators, histone modifiers, or lncRNAs to interact effectively with target loci, due to their loss of DNA binding specificity and low intracellular concentrations [47, 49]. This underscores the value of using dCas9-based CAPTURE, as illustrated by [47], which can identify interacting proteins, RNAs and DNAs, ultimately providing insights into parasitism gene regulation models. Remarkably, regarding effectors loci, sgRNAs could be designed to target specific groups of effectors, provided that conserved sequences exist among their promoters. Beyond candidate identification, the methodology offers the potential to unravel the signalling networks controlling effector gene expression.

However, our approach does not determine whether the chromatin-binding proteins identified originated from gland cells or other cell types. It is conceivable that these proteins may function as positive regulators in gland cells and as negative regulators in other cells of the animal, and so subsequent *in situ* hybridisation to the putative interactors is required [50].

## Conclusion

In this study, we successfully developed and optimized an *in vitro* dCas9-based CAPTURE method based on the CRISPR-dCas9 system, specifically tailored for *Meloidogyne incognita*. Despite technical challenges in nematode material preparation, we established a reproducible protocol for chromatin isolation and fragmentation, enabling various dCas9-based CAPTURE assays. This represents a significant advancement for precise genomic analysis in non-transformable PPN. Our identification of BANF1 as a protein candidate, which is linked to chromatin remodelling complexes like SWI/SNF, suggests its potential role in the epigenetic regulation of parasitism. These findings underscore the importance of transcriptional and epigenetic mechanisms in the precise regulation of parasitism gene expression. However, the exact roles of these chromatin-binding proteins require further functional validation, such as through RNAi-based approaches and *in situ* hybridisation. Additionally, the flexibility of our approach highlights its broader potential. The dCas9-based CAPTURE method could be extended to target specific groups of effector genes by designing sgRNAs that recognize conserved sequences in their promoters. This opens up new avenues for understanding the signalling networks that regulate effector gene expression, which could be crucial for advancing our knowledge of parasitism in *M. incognita*. Future research should focus on expanding this method to additional effector loci, offering insights that could ultimately inform novel strategies for pest management.

## Material and Methods

### Nematode collection

*Meloidogyne incognita* race Morelos acquired from Laboratory of Nematology of Wageningen University & Research was used in all experiments. The nematode culture and maintenance were conducted on greenhouse tomato roots (*Solanum lycopersicum* cv. Roma) growing in a mixture of substrate/sand (ratio 2:1). Infected plants were watered daily and maintained at approximately 23°C all the year. Egg nematodes were collected every (8-11) weeks for reinoculation (depending on the period during the year). The nematode extraction and hatching were as described in [51]. Worms were freshly collected every two days, then left to settle at room temperature and subsequently cleaned with 500µl HBSS buffer after centrifugation at 13, 000 rpm for 1 min. After removing the homemade Hank’s Balanced Salt solution (HBSS) buffer (5mM KCl, 0,4mM KH2PO4, 4mM NaHCO3, 50mM NaCl, 0,3mM Na2HPO4, 5,5mM Glucose (filter 0,2μm)) buffer, nematode aliquots were stored at -80°C for several weeks, termed hereafter frozen nematodes. All of the experimentations have been conducted from aliquots of 30,000 ppJ2 or eggs.

### dCas9-based CAPTURE procedure based on *in vitro* CRISPR-dCas9 system Standard disruptions of worms with different grinding devices

Thirty thousand larvae or eggs were resuspended in 150 µl cold HBSS buffer supplemented with 100 X freshly prepared cocktail of protease inhibitors (Roche, catalog # 11873580001) and transferred into distinct handling devices for mechanical disruption of worms: 1) 2-ml Dounce Tissue Grinder (Kimble kontes®) combined with either the large-clearance pestle A (0.063 - 0.14mm - loose pestle) or the small-clearance pestle B (0.0127 - 0.0635mm - tight pestle), 2) the needle/syringe system with 29-gauge needle (29G, 0.33mm x 13mm) and 3) Balch homogenizer using different combinations of ball bearing sizes and syringe pass numbers i) 18 µm to 14 µm; ii) 14 to 6 µm and iii) 14 µm to 4 µm ball bearing for 25 passes for each condition. Disruption with the Balch system was conducted in a hypotonic buffer (10 mM HEPES pH7.6, 10 mM KCl, 1,5 mM EDTA, 250 mM sucrose). As reference, the use of Balch for complete disruption of L4 larvae *C. elegans* (twice longer than ppJ2 *M. incognita*) required 25 passes with an 18µm clearance ball bearing followed by 25 passes with a 12 µm -clearance as described in [29, 30].

To monitor the progress of the mechanical disruption, a time course was performed at different times and worms were visually examined by optical microscope. We picked up 5 μl of larval suspension, and diluted 10-fold for examination under an optical microscope. The homogeneous nematode extracts were then subjected to chemical disruption. Following steps were conducted at 4°C or in a cold chamber room. Hereafter, all buffers of chromatin preparation kit (Diagenode Chromatin EasyShear kit - Low SDS (C01020013)) were prepared on experimental day and freshly supplemented with a cocktail of protease inhibitors (Roche, catalog # 11873580001). First, the pellet was resuspended in 100 µl cold lysis IL1b buffer and incubated for 10 min (rotating wheel, 21 rpm in a cold chamber room), then centrifuged for 5 min at 2700 rpm. In similar experimental conditions, the pellet was incubated with 100µl cold lysis IL2 buffer and then with 100 µl complete Shearing IS1 buffer. Samples were lastly centrifuged at 20, 000 x g for 10 min. Samples were stored at 4°C for subsequent quality and quantity assessments (including agarose gel electrophoresis, bioanalyzer, qubit for DNA or SDS-PAGE and western blotting for proteins).

### Sonication treatment for chromatin fragmentation

Cell pellets were rinsed in 100 µl iS1 buffer (rotating wheel, 21 rpm in a cold chamber room) for 10 min, subsequently transferred into 1.5 ml TPX microtubes (#M-50001, Diagenode). Sonication was performed on the Diagenode Bioruptor PLUS instrument (#B01020004) using standard settings: high (H) power input, 30s ON/30s OFF pulses, +4°C water bath but no floating ice. Sonication conditions were specified in the captions of figures. Immediately after sonication, chromatin was cleared in a hard spin at 16, 000 x g for 10 min at 4°C. Supernatants (SN) containing the sheared chromatin were pooled in batch into a new 1,5-ml eppendorf tube, and then aliquoted for further chromatin experiments (DNA purification, SDS-PAGE and Western blotting analyses). Samples can be stored and frozen at -20°C and -80°C for later use.

### Single-step sonication procedure for simultaneous cell disruption and chromatin shearing

Beforehand, worms were subjected to the cross-linking reaction. Aliquots of HBSS- cleaned 30,000 ppJ2 or eggs stored at -80°C were thawed and shortly stabilized at room temperature before adding the crosslinker agent (i.e formaldehyde), because kinetics and efficiency of formaldehyde-mediated cross-linking are temperature-dependent [52]. To improve cross-linking reproducibility, a single-use 37% formaldehyde ampule was used at experimental day (e.g Pierce) and all cross-linking steps were conducted at room temperature in the rotating wheel (21 rpm). Hereafter, all buffers were supplemented freshly prepared cocktail of protease inhibitors (Roche, catalog # 11873580001). First, worms were resuspended in 500 µl of fixative buffer. Crosslinker concentration and time incubation were specified in the caption of figures. Cross-linked worms were quenched with 750 mM Tris-HCl for 5 min [53]. Hereafter nematode extracts were maintained at 4°C and washed twice in 500 µl HBSS cold buffer. Worm pellets were collected by centrifugation at 2,700 rpm for 5 min at 4°C. After discarding the supernatant, the worm pellet was resuspended in 100 µl cold iS1 buffer and incubated for 10 min on a rotating wheel. The cross-linked animals were then sonicated with a Bioruptor PLUS instrument to disrupt worms and simultaneously shear chromatin to desired size for R-ChIP assays. The sonication parameters were as follows: high (H) power input, i.e 20 cycles of 90s ON/15s OFF pulses - mixing gently tubes every 10 cycles - and 10 additional cycles of 30s ON/15s OFF pulses, +4°C water bath but no floating ice. This instrument allows to sonicate up to 6 samples per run. The sonicated chromatin was collected from worm lysates after centrifugation at 16,000 x g for 10 min at 4°C. Visual inspection of the pellet after centrifugation was conducted by optical microscope to evaluate the impact of sonication on worm disruption. Supernatant aliquots were harvested and used for further chromatin studies experiments.

### Reverse crosslinking and DNA recovery

To evaluate the effect of sonication procedure on chromatin shearing, we conducted reverse DNA crosslinking and purification. As described in [53], 20 µl of sheared chromatin was de-cross-linked with the addition of 95 µl TE/SDS buffer (10 mM Tris pH8 1 mM EDTA 1% SDS) and incubated overnight at 65°C in a thermoshaker (800 rpm). Day after, samples were treated with 2,5 µl RNAseA (10 mg/ml) for 30 min at 37°C followed by the addition of 20 µl proteinase K (10 µg/µl) and incubation at 55°C for 2 h. Samples were then cleaned up with MicroChIP DiaPure columns kit (#C03040001, Diagenode) and eluted with 8 µl nuclease free water, following the manufacturer’s instructions. Fragmented DNA samples were visualized on agarose gel electrophoresis 1,5% by loading samples at 250 ng per lane and on an Agilent 2100 Bioanalyzer High Sensitivity DNA Chip (#G2938-68000) by following the manufacturer’s recommendation.

### Analysis of proteins SDS-PAGE silver-stained and Western blotting

The collected pellet and all supernatant fractions were resuspended in Laemmli urea 1X or 2X (126 mM Trsi-HCl ph6.8, 8M Urea, 20% Glycerol, 4% SDS, 0,02% bromophenol blue) respectively, then boiled at 80°C for 8 min. Aliquots with a loading amount at 500 J2 were used for SDS-PAGE analysis (Mini-PROTEAN TGX precast protein gels 4-15%, #4561086) and stained using the Pierce Silver Stain kit (Thermo Scientific ™) by following the manufacturer’s protocol. Similarly, aliquots from each fraction were separated on SDS-PAGE 4-15% gradient gel and transferred to a PVDF membrane with western blot settings: 14V/1.3 mM 7 min on Biorad instrument. Membranes were rinsed twice in TBS with 0,1% Tween-20 for 5 min and then blocked overnight with 5% milk at 4°C O/N under agitation. Next, blots were incubated with the primary antibody rabbit polyclonal anti-histone H3 antibody (#ab1791) diluted 1:2500 and then with the secondary, the goat anti-rabbit IgG antibody conjugated to HRP at 1: 10,000 dilution, both for 1 hour at room temperature in 3% milk TBS buffer with 0,1% Tween-20. Signals were then developed for 3 min using chemiluminescent kit (Clarity ™ Western ECL substrate, #1705062). Immunoblots were imaged using Biorad ChemiDoc XRS+ Gel Imaging System.

### *in vitro* dCas9-based CAPTURE

Protocol for the in vitro dCas9-based CAPTURE assay was adapted from the experimental procedure realized in [35, 36, 54].

### *in vitro* dCas9-based CAPTURE using genomic DNA as bait

The procedure of *in vitro* dCas9-based CAPTURE from gDNA of 50,000 ppJ2 *M. incognita* were as follows : **Genomic DNA extraction** - Worms were ground into 50 µl of lysis buffer (0,1M Tris pH8.0, 0,5M NaCl, 50 mM EDTA and 1% SDS) with pestle for 30 sec. and then added 450 µl of lysis buffer, 4 µl proteinase K (20 mg/ml) for an incubation of 24 hours at 55°C in thermomix. The worm lysate was next treated with 10 µl RNAse A (10 mg/ml) for 30 min at 37°C. **Genomic DNA precipitation** - gDNA was precipitated by adding Na-Ac buffer (Ammonium acetate 2.5 M, sodium acetate 0,3M) volume equivalent, then completed with 0,7 volume of isopropanol at room temperature and mixed. The worm solution was centrifuged at 15, 000 x g for 30 min at 4°C . The pellet was then washed with 70% Ethanol volume equivalent, centrifuged at 15, 000 x g for 15 min at 4°C and dried for 10 min at room temperature. DNA extract was resuspended with 8 µl nuclease free water for 2 hours at 55°C. **Genomic DNA sonication and purification** - Cleaned gDNA was fragmented in the cold iS1 buffer with a final volume of 100 µl. sonication parameters were as follows: high (H) power input, i.e 5 cycles of 30s ON/ 30s OFF pulses, +4°C water bath but no floating ice. The average length of DNA fragments was approximately 400-700 bp. After sonication, sheared gDNA was purified with MicroChIP DiaPure columns kit (#C03040001, Diagenode) and eluted with 8 µl nuclease free water with a final concentration around 6µg (ratio 260/280 nm 1.85). **Precleaning of magnetic beads** - Thirty microliters of magnetic beads (Thermo Scientific ™ Pierce ™ Streptavidin magnetic beads, #PI88816) were added to 1,5- mL eppendorf tube, harvested and cleaned twice with 500 µl of washing buffer PBST (1X PBS pH7.4, 10% Tween- 20) using the magnetic frame for 3 min on ice. Lastly magnetic beads were resuspended with 500 µl PBST-B buffer (completed with 0.1% BSA). **Hybridizing dCas9-RNP complex to magnetic beads** - According to literature curation, the RNP dCas9-sgRNA complex was prepared at a molar ratio 1:5 for hybridizing 1 µg of sheared gDNA. Then 1,56 µg of dCas9 recombinant protein [36] was incubated with pre-cleaned magnetic beads in the PBST-B buffer for 2-3 hours (rotation wheel, in cold chamber) and placed on ice for 3 min with the magnetic frame to discard the supernatant. Next, magnetic beads complex was washed twice and resuspended in 100 µl in vitro CRISPR buffer (20 mM Hepes pH7,1, 150 mM KCl, 0,1 mM EDTA, 10 mM MgCl2 add the same day 0,5 mM DTT 1X Protease inhibitors EDTA free (Roche) and 0,85 µg of each sgRNA was added to dCas9/beads mix for 10 min of incubation at 37°C. **gDNA hybridization and affinity isolation using the *in vitro* dCas9-CRISPR system** - One microgram of cleaned gDNA was first diluted 10-fold into in vitro CRISPR buffer (exposure to SDS present in iS1 buffer could impact on the dCas9-RNP function and stability). The gDNA was conjugated to the mix previously prepared and incubated with 40U/ µl RNAsin (Promega) at 37°C for 20 min under agitation at 650 rpm in Thermomix. The magnetic beads were washed sequentially with the following buffers using the magnetic frame: washing fourth with 1-mL low salt buffer (20 mM Tris pH 8.0, 150 mM NaCl, 2 mM EDTA, 0,1 % Triton, 0,03% SDS supplemented with 5U/ml RNAsin (invitrogen) and 1X Protease inhibitor cocktail), washing once with 1-mL TBS-NP40 (50 mM Tris pH 7.5, 150 mM NaCl, 0,5% NP-40), washing thrice with 1-mL TE (10 mM Tris pH7.5, 1 mM EDTA). Then remove the supernatant and eluted with 100 µl SDS-ChIP buffer (20 mM Tris pH 8.0, 2mM EDTA, 150 mM NaCl, 0,1 % SDS, 1% Triton X100 supplemented with 1X Protease inhibitor cocktail) for overnight incubation at 70°C into thermocycler. The day after, samples were cooled down at room temperature and completed with 100 µl TE buffer. The captured gDNA fragments were then purified with MicroChIP DiaPure columns kit (#C03040001, Diagenode) and used as a DNA template to determine the hybridization efficiency assay. The gene promoter for MiActin was used as the internal reference. The truncated 6F06 promoter was PCR amplified using primers and the DNA from samples with and without dCas9-RNP as DNA templates. The primers used in the study are shown in Supplementary Table S2.

### *in vitro* dCas9-based CAPTURE using native chromatin as bait

The preparation of biological material, crosslinking and sonication were performed as described above. Crosslinking and sonication parameters were specified in the captions of their respective figures. One sample test corresponded to 5 µg cross-linked and sheared chromatin obtained from 30, 000 ppJ2. After centrifugation, the input fragmented chromatin (peaking at 100-500 bp) was immunoprecipitated using dCas9-biotin system as described above. The captured fractions were used for subsequent experiments (Bioanalyzer, mass spectrometry, SDS-PAGE analysis).

#### MS-based proteomic analyses

Proteins were solubilized in Laemmli buffer and stacked in the top of a 4-12% NuPAGE gel (Invitrogen), stained with Coomassie blue R-250 (Bio-Rad) before in-gel digestion using modified trypsin (Promega, sequencing grade) as previously described [55]. The resulting peptides were analyzed by online nanoliquid chromatography coupled to MS/MS (Ultimate 3000 RSLCnano and Orbitrap Exploris, Thermo Fisher Scientific) using a 60-min gradient. For this purpose, the peptides were sampled on a precolumn (300 μm x 5 mm PepMap C18, Thermo Scientific) and separated in a 75 μm x 250 mm C18 column (Aurora Generation 3, 1.7μm, IonOpticks). The MS and MS/MS data were acquired using Xcalibur (Thermo Fisher Scientific).

Peptides and proteins were identified by Mascot (version 2.8.0, Matrix Science) through concomitant searches against homemade databases containing the predicted protein sequences of *M. incognita* and of classical contaminant proteins found in proteomic analyses (keratins, trypsin…). Trypsin/P was chosen as the enzyme and two missed cleavages were allowed. Precursor and fragment mass error tolerances were set at respectively at 10 and 20 ppm. Peptide modifications allowed during the search were: Carbamidomethyl (C, fixed), Acetyl (Protein N-term, variable) and Oxidation (M, variable). The Proline software [56] (version 2.3.3) was used for the compilation, grouping, and filtering of the results (conservation of rank 1 peptides, peptide length ≥ 6 amino acids, false discovery rate of peptide-spectrum-match identifications < 1% [57], and minimum of one specific peptide per identified protein group). Proline was then used to perform a weighted spectral counting (WSC)-based comparison of the samples. Proteins were considered enriched in a sample compared to another if they were identified by at least 3 WSC and identified only in one sample or enriched at least 5 times in a sample on the basis of WSC.

### Cas9 digestion assay

To evaluate the efficiency and specificity of sgRNAs, DNA cleavage assay was conducted on PCR DNA targets of 6F06 gene locus (Minc3s08477g42291). Target region was amplified from *M. incognita* genomic DNA with specific primers (Table S2) using the Phusion High-Fidelity PCR kit (New England Biolabs) prior to analysis by gel electrophoresis and then purified using a purification kit (Monarch). The Cas9 cleavage reaction consisted of adding in thermocycler: 260 nM sgRNA, 260 nM Cas9 protein (CAS9PROT, Sigma) 10X NEB3.1 for 10 min at 25°C, then added 26 nM purified PCR product of interest for 1 hour at 37°C with final molar ratio 10:10:1. Afterwards, Cas9 was heat inactivated for 20 min at 80°C. The results of cleaving effector gene fragments were assessed by electrophoresis on 2% agarose gel.

## Supplementary data

**Supplementary Figure S1. Optimization of homogenization methods for the PPN *Meloidogyne incognita*. (A)** Photos of the Balch homogenizer (left) and Dounce homogenizer (right). **(B)** Approximately 30,000 second-stage larvae (J2) were resuspended in homemade Hank’s Balanced Salt solution (HBSS) buffer and gently disrupted on ice to generate a relative homogenate. Worm lysates were processed using various mechanical homogenization techniques: Dounce homogenizer with loose (A) or tight (B) pestle, Balch homogenizer using an 4- to 14 µm clearance ball bearing and with different combinations of syringe pass numbers, and Bioruptor device under different time and cycle conditions. All homogenization steps were performed on ice or at 4°C to maintain sample integrity. While these methods were effective in grinding nematode cells to subcellular components, none achieved complete homogenization. Further validation steps, including nuclear extract preparation, chromatin proteome analysis, and DNA isolation, were performed to ensure sample quality and compatibility for downstream analyses. Scale bar: 50µm

**Supplementary Figure S2. Chromatin isolation from *M. incognita* larvae using the dounce homogenizer with the pestle B. (A)** Schematic outlining the steps for chromatin isolation from pre-parasitic juveniles (ppJ2), using the dounce system (pestle B). Starting material consists of 30,000 ppJ2 incubated in root extract for 2 hours, followed by flash-freezing in liquid nitrogen and storage at -80°C. Samples from each step were separated into supernatant (**TP, SOL, L1, L2, SN**) and pellet (or nuclear crude extract, P) fractions. **(B)** Each lane contains protein extract from 500 ppJ2. Silver-stained SDS-PAGE (4-15%) illustrates the protein profile at key stages of chromatin isolation (upper panel). Immunoblotting for histone H3 (bottom panel) highlights chromatin enrichment. Exposure time for chemiluminescence is indicated in seconds (s)). Histones presence (≈16 kDA) is marked with red stars, confirmed by the anti-H3 antibody.

**Supplementary Figure S3. Protein extracts after homogenization and sonication steps from *M. incognita* eggs. (A)** Pipeline for chromatin isolation and shearing process using the Diagenode Bioruptor® as the sonication device. (B) Frozen eggs were directly sonicated, followed by centrifugation. Aliquots of protein extracts, comprising supernatant **(SN)** and pellet **(P)** fractions, were collected and separated by SDS-PAGE (4-15%), then visualized by silver staining. The presence of histones H3 (≈16 kDa) was not detected by immunoblotting using an anti-H3 antibody.

**Supplementary Figure S4. Effect of crosslinking conditions on chromatin-protein isolation and DNA shearing in *M. incognita* eggs. (A)** Pipeline for chromatin preparation following crosslinking of eggs, utilizing the Diagenode Bioruptor® as the sonication device. Frozen eggs were crosslinked using formaldehyde, then centrifuged before formaldehyde quenching (Tris-HCl, 5 min). Samples were subsequently centrifuged again, and pellets were washed with a physiological buffer. Crosslinked eggs were directly sonicated, followed by centrifugation, and their sheared chromatin profile was analyzed using silver-stained SDS-PAGE and agarose gel electrophoresis. **(B)** Aliquots of protein extracts, comprising supernatant **(SN)** and pellet **(P)** fractions, were collected, separated by SDS-PAGE (4-15%), then visualized by silver staining. The presence of histones was not observed by immunoblotting with histone H3 antibody. **(C)** Chromatin samples generated in **(B)** were analyzed by SYBRSafe DNA electrophoresis in 2% agarose gel. Fixation at 1 % formaldehyde for 10 min presents optimal time and concentration for shearing chromatin, where most DNA fragments were distributed from 100 to 400 bp.

**Supplementary Figure S6. *in vitro* Cas9 assay to assess the cleavage efficiency from chosen sgRNAs targeting PCR-amplified 6F06 locus. (A)** Map of the 6F06 effector genomic locus (Minc3s08477g42291), showing the regions targeted by the recombinant Cas9/sgRNA complexes. The promoter region contains Mel-DOG boxes, existing in two variants (blue and turquoise arrows). The 6F06 promoter region is targeted with two ribonucleoprotein (RNP) complexes, each composed of the active form of Cas9 and a specific sgRNA (sg1 and sg2), with their positions are indicated. PCR amplification was performed using flanking primers (grey arrows) to generate the DNA template for the assay **(B)** Representative agarose gel showing results from in vitro digestion assays testing the cleavage efficiency of two sgRNAs targeting the PCR-amplified 6F06 locus. Purified PCR matrices incubated with recombinant Cas9 protein lacking a specific sgRNA (-) showed no cleavage, while the addition of sgRNA/Cas9 complexes led to moderate cleavage of the target PCR product. M = 1 kb DNA ladder.

**Supplementary table S1.** Sequences of single guide RNAs

**Supplementary table S2.** Primers used in this study

**Supplementary table S3.** MS-based comparative analysis of proteins eluted from +RNP and - RNP samples

## Acknowledgments

Caroline Bournaud would like to thank Cecile Bedet (ENS Lyon) and Rahim Hassanaly-Goulamhoussen (INRAE Sophia Antipolis) for helpful discussions for homogenization devices applied in this study. Caroline Bournaud is thankful to Elise Lacroix from the greenhouse platform of the University Claude Bernard Lyon1 (UCBL1) for her careful management of tomato plants. *Melodoigyne incognita* eggs were kindly provided by Casper C. van Schaik from Laboratory of Nematology of Wageningen University and Research. We thank François-Xavier Gillet for helpful advice in performing *in vitro* dCas9-based CAPTURE experiments and critical reading on the manuscript, and Nathalie Poussereau for fruitful comments during the drafting of the manuscript. Caroline Bournaud received funding from the European Union’s Horizon 2020 Research and innovation programme under the Marie Skłodowska-Curie Grant Agreement No 833420. Work on plant-parasitic nematodes at the University of Cambridge is supported by DEFRA licence 125034/359149/3, and funded by BBSRC grants BB/R011311/1, BB/S006397/1, BB/X006352/1, and BB/Y513246/1 a Leverhulme grant RPG-2023-001, and a UKRI Frontier Research Grant EP/X024008/1. The proteomic experiments were partially supported by Agence Nationale de la Recherche under projects ProFI (Proteomics French Infrastructure, ANR-10-INBS-08) and GRAL, a program from the Chemistry Biology Health (CBH) Graduate School of University Grenoble Alpes (ANR-17-EURE-0003).

## Declarations

● **Ethics approval and consent to participate**

Not applicable.

**● Consent for publication**

Not applicable

● **Availability of data and materials**
● Competing interests

The authors declare no competing interests.

## Authors’ contributions

CB and SEvDA conceptualized and designed the research study, CB performed the experiments and optimizations. SEVDA and EGJD performed bioinformatic analyses. AT and YC performed MS-based proteomic experiments. SEVDA is the principal investigator of the research grant. CB wrote the manuscript, assisted by all the authors.

