## Supplementary S1 for "DNA-protein interactions in *Meloidogyne incognita*"

### Slide 1
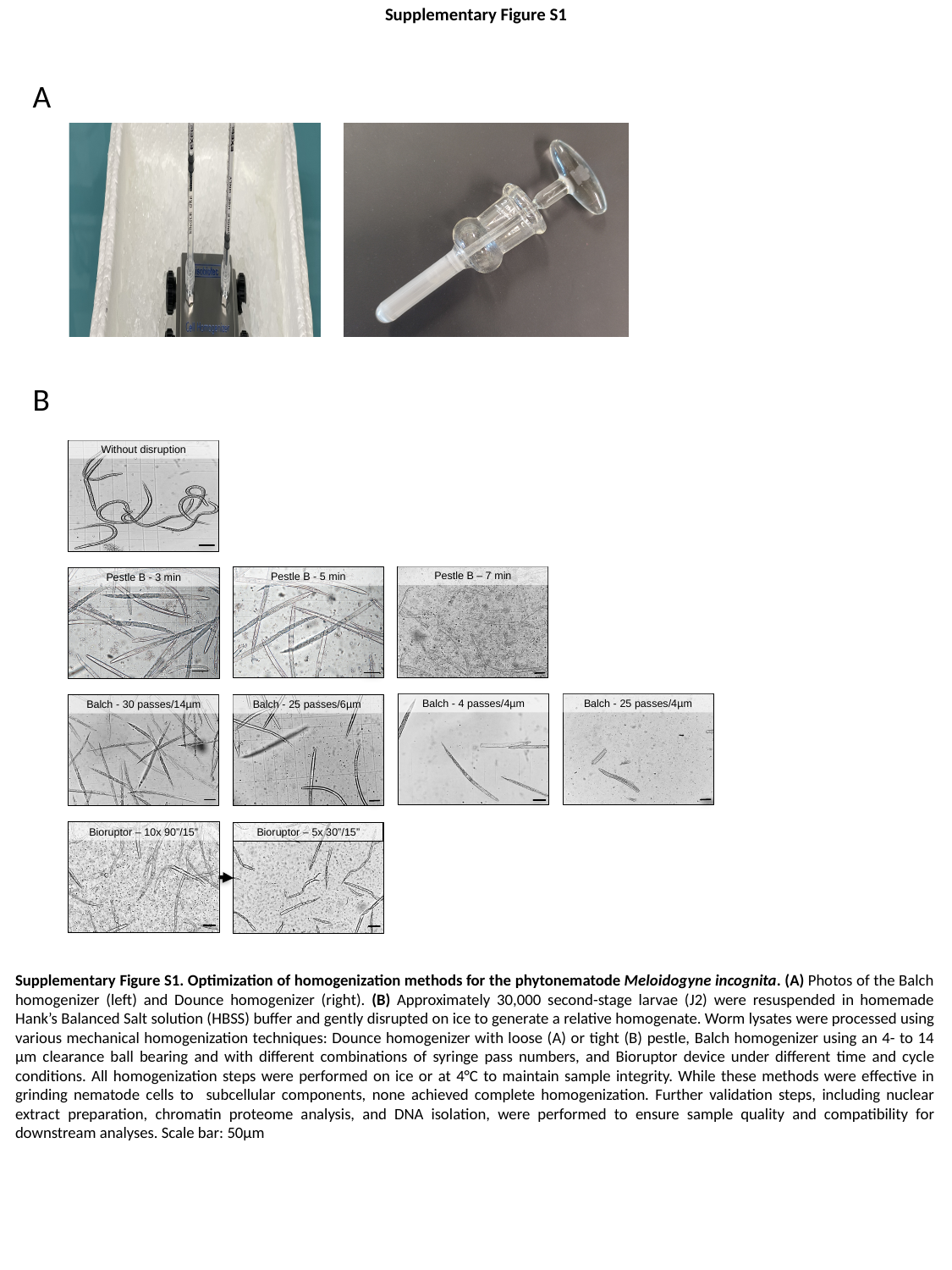

Supplementary Figure S1
A
B
Without disruption
Pestle B – 7 min
Pestle B - 5 min
Pestle B - 3 min
Balch - 4 passes/4µm
Balch - 25 passes/4µm
Balch - 30 passes/14µm
Balch - 25 passes/6µm
Bioruptor – 10x 90”/15”
Bioruptor – 5x 30”/15”
Supplementary Figure S1. Optimization of homogenization methods for the phytonematode Meloidogyne incognita. (A) Photos of the Balch homogenizer (left) and Dounce homogenizer (right). (B) Approximately 30,000 second-stage larvae (J2) were resuspended in homemade Hank’s Balanced Salt solution (HBSS) buffer and gently disrupted on ice to generate a relative homogenate. Worm lysates were processed using various mechanical homogenization techniques: Dounce homogenizer with loose (A) or tight (B) pestle, Balch homogenizer using an 4- to 14 µm clearance ball bearing and with different combinations of syringe pass numbers, and Bioruptor device under different time and cycle conditions. All homogenization steps were performed on ice or at 4°C to maintain sample integrity. While these methods were effective in grinding nematode cells to subcellular components, none achieved complete homogenization. Further validation steps, including nuclear extract preparation, chromatin proteome analysis, and DNA isolation, were performed to ensure sample quality and compatibility for downstream analyses. Scale bar: 50µm
