## Supplementary S2 for "DNA-protein interactions in *Meloidogyne incognita*"

### Slide 1
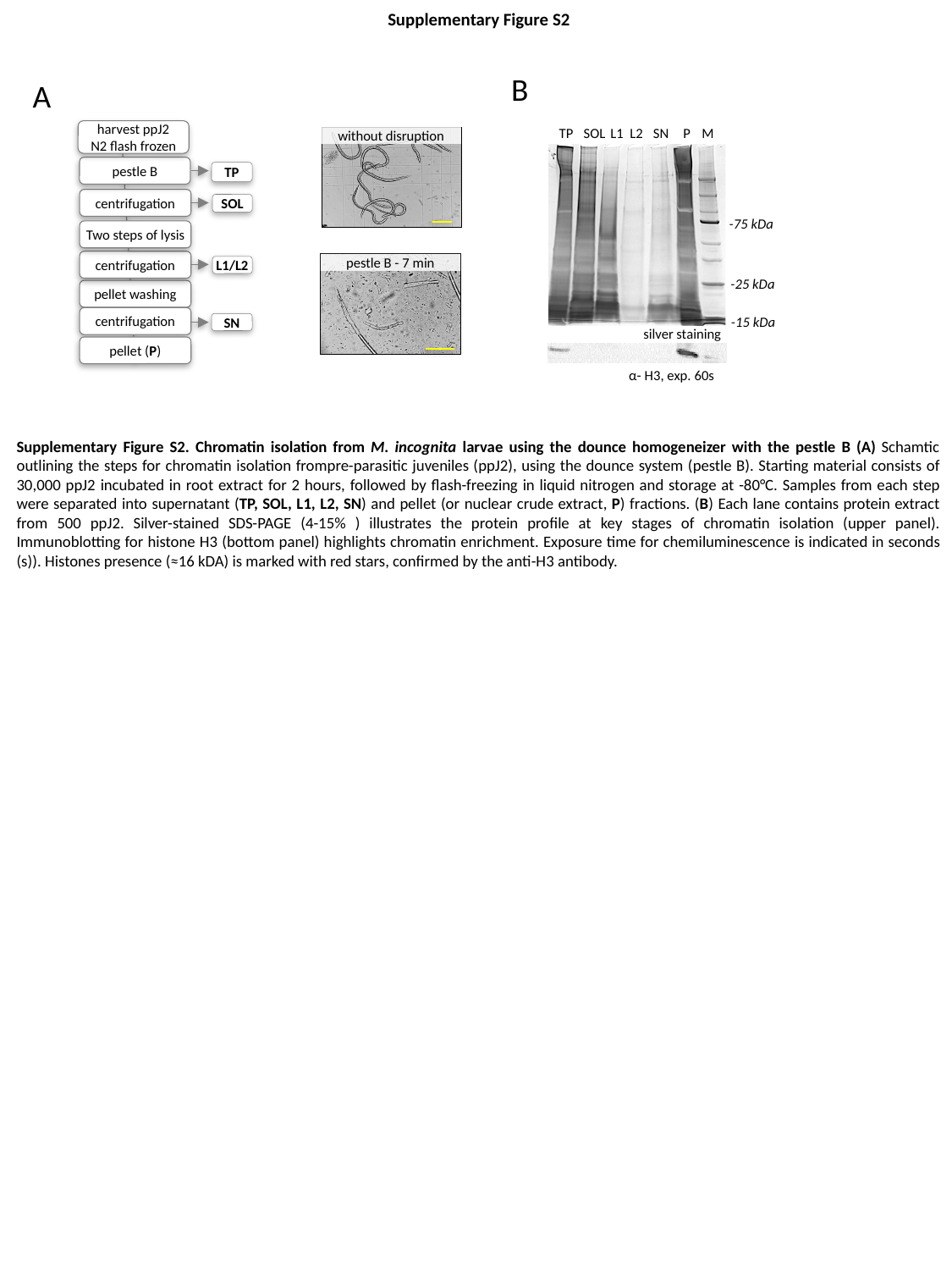

Supplementary Figure S2
B
A
harvest ppJ2
N2 flash frozen
pestle B
TP
centrifugation
SOL
Two steps of lysis
centrifugation
L1/L2
pellet washing
centrifugation
SN
pellet (P)
| TP | SOL | L1 L2 | SN | P | M |
| --- | --- | --- | --- | --- | --- |
without disruption
-75 kDa
-25 kDa
silver staining
-15 kDa
silver staining
α- H3, exp. 60s
pestle B - 7 min
Supplementary Figure S2. Chromatin isolation from M. incognita larvae using the dounce homogeneizer with the pestle B (A) Schamtic outlining the steps for chromatin isolation frompre-parasitic juveniles (ppJ2), using the dounce system (pestle B). Starting material consists of 30,000 ppJ2 incubated in root extract for 2 hours, followed by flash-freezing in liquid nitrogen and storage at -80°C. Samples from each step were separated into supernatant (TP, SOL, L1, L2, SN) and pellet (or nuclear crude extract, P) fractions. (B) Each lane contains protein extract from 500 ppJ2. Silver-stained SDS-PAGE (4-15% ) illustrates the protein profile at key stages of chromatin isolation (upper panel). Immunoblotting for histone H3 (bottom panel) highlights chromatin enrichment. Exposure time for chemiluminescence is indicated in seconds (s)). Histones presence (≈16 kDA) is marked with red stars, confirmed by the anti-H3 antibody.
