## Supplementary S3 for "DNA-protein interactions in *Meloidogyne incognita*"

### Slide 1
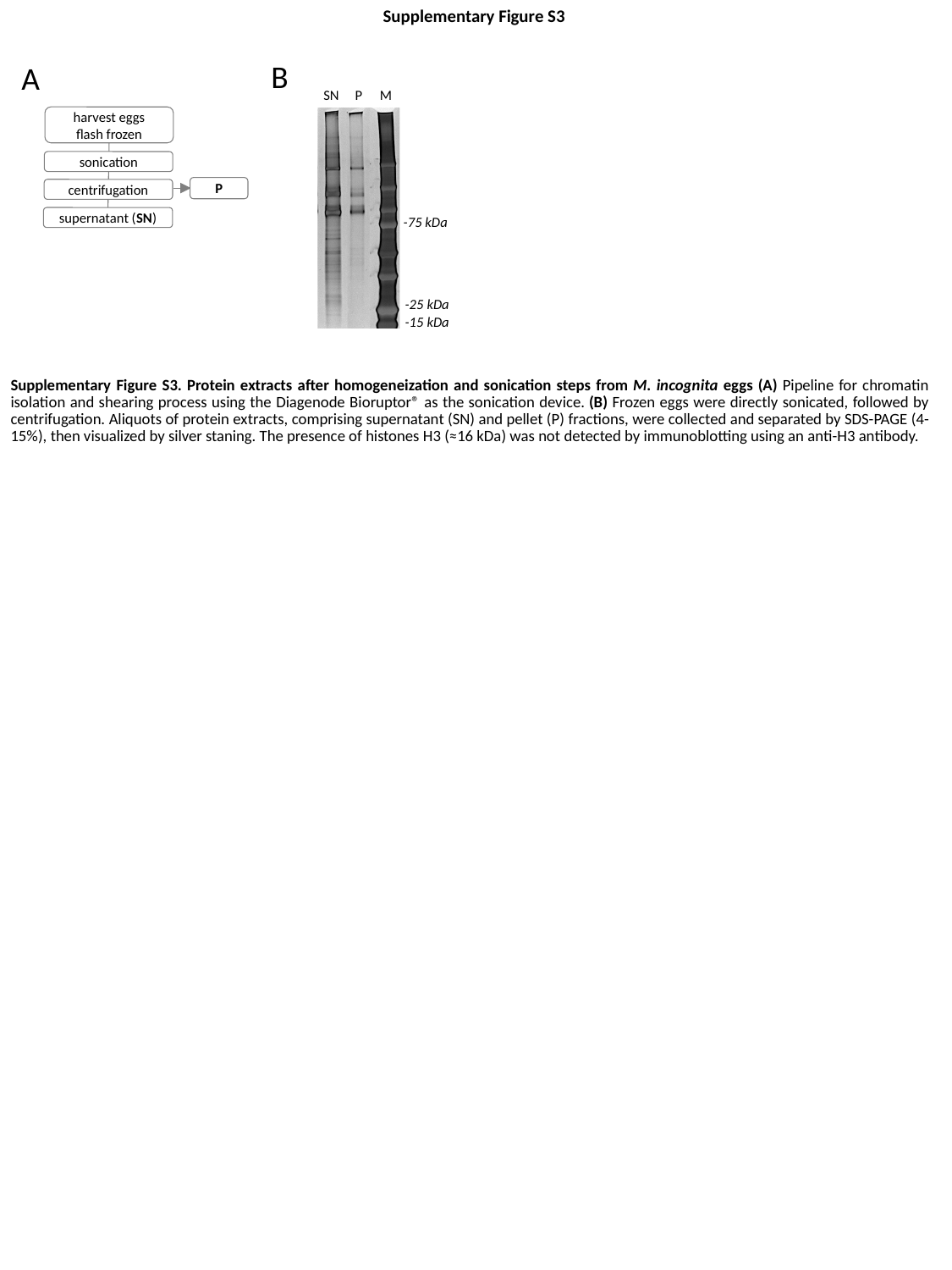

Supplementary Figure S3
B
A
| SN | P | M |
| --- | --- | --- |
harvest eggs
flash frozen
sonication
P
centrifugation
-75 kDa
supernatant (SN)
-25 kDa
-15 kDa
Supplementary Figure S3. Protein extracts after homogeneization and sonication steps from M. incognita eggs (A) Pipeline for chromatin isolation and shearing process using the Diagenode Bioruptor® as the sonication device. (B) Frozen eggs were directly sonicated, followed by centrifugation. Aliquots of protein extracts, comprising supernatant (SN) and pellet (P) fractions, were collected and separated by SDS-PAGE (4-15%), then visualized by silver staning. The presence of histones H3 (≈16 kDa) was not detected by immunoblotting using an anti-H3 antibody.
