## Supplementary S4 for "DNA-protein interactions in *Meloidogyne incognita*"

### Slide 1
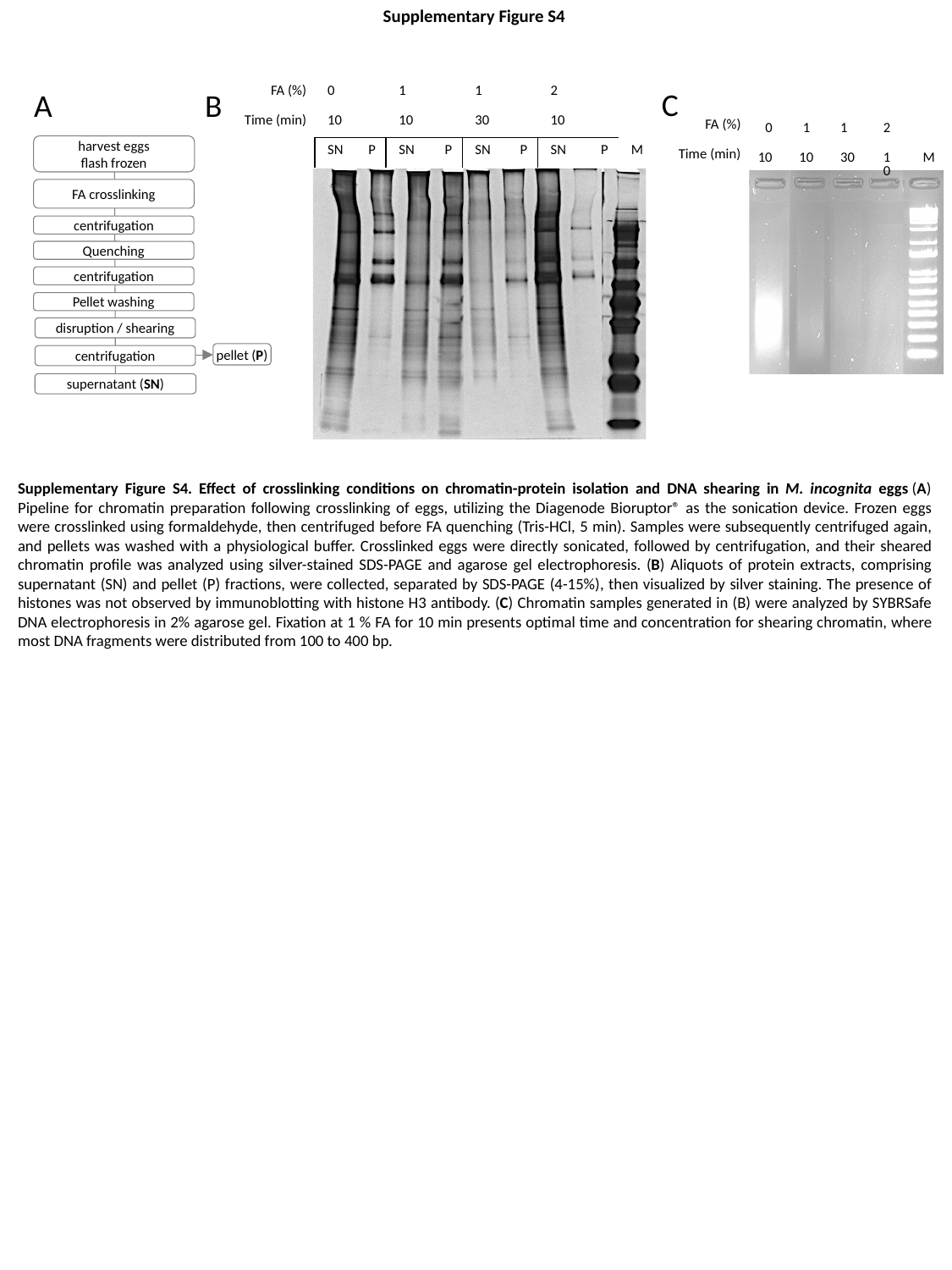

Supplementary Figure S4
| 0 | | 1 | | 1 | | 2 | | |
| --- | --- | --- | --- | --- | --- | --- | --- | --- |
| 10 | | 10 | | 30 | | 10 | | |
| SN | P | SN | P | SN | P | SN | P | M |
C
| FA (%) |
| --- |
| Time (min) |
A
B
| FA (%) |
| --- |
| Time (min) |
| 0 | 1 | 1 | 2 | |
| --- | --- | --- | --- | --- |
| 10 | 10 | 30 | 10 | M |
harvest eggs
flash frozen
FA crosslinking
centrifugation
Quenching
centrifugation
Pellet washing
disruption / shearing
pellet (P)
centrifugation
supernatant (SN)
Supplementary Figure S4. Effect of crosslinking conditions on chromatin-protein isolation and DNA shearing in M. incognita eggs (A) Pipeline for chromatin preparation following crosslinking of eggs, utilizing the Diagenode Bioruptor® as the sonication device. Frozen eggs were crosslinked using formaldehyde, then centrifuged before FA quenching (Tris-HCl, 5 min). Samples were subsequently centrifuged again, and pellets was washed with a physiological buffer. Crosslinked eggs were directly sonicated, followed by centrifugation, and their sheared chromatin profile was analyzed using silver-stained SDS-PAGE and agarose gel electrophoresis. (B) Aliquots of protein extracts, comprising supernatant (SN) and pellet (P) fractions, were collected, separated by SDS-PAGE (4-15%), then visualized by silver staining. The presence of histones was not observed by immunoblotting with histone H3 antibody. (C) Chromatin samples generated in (B) were analyzed by SYBRSafe DNA electrophoresis in 2% agarose gel. Fixation at 1 % FA for 10 min presents optimal time and concentration for shearing chromatin, where most DNA fragments were distributed from 100 to 400 bp.
