## Supplementary S5 for "DNA-protein interactions in *Meloidogyne incognita*"

### Slide 1
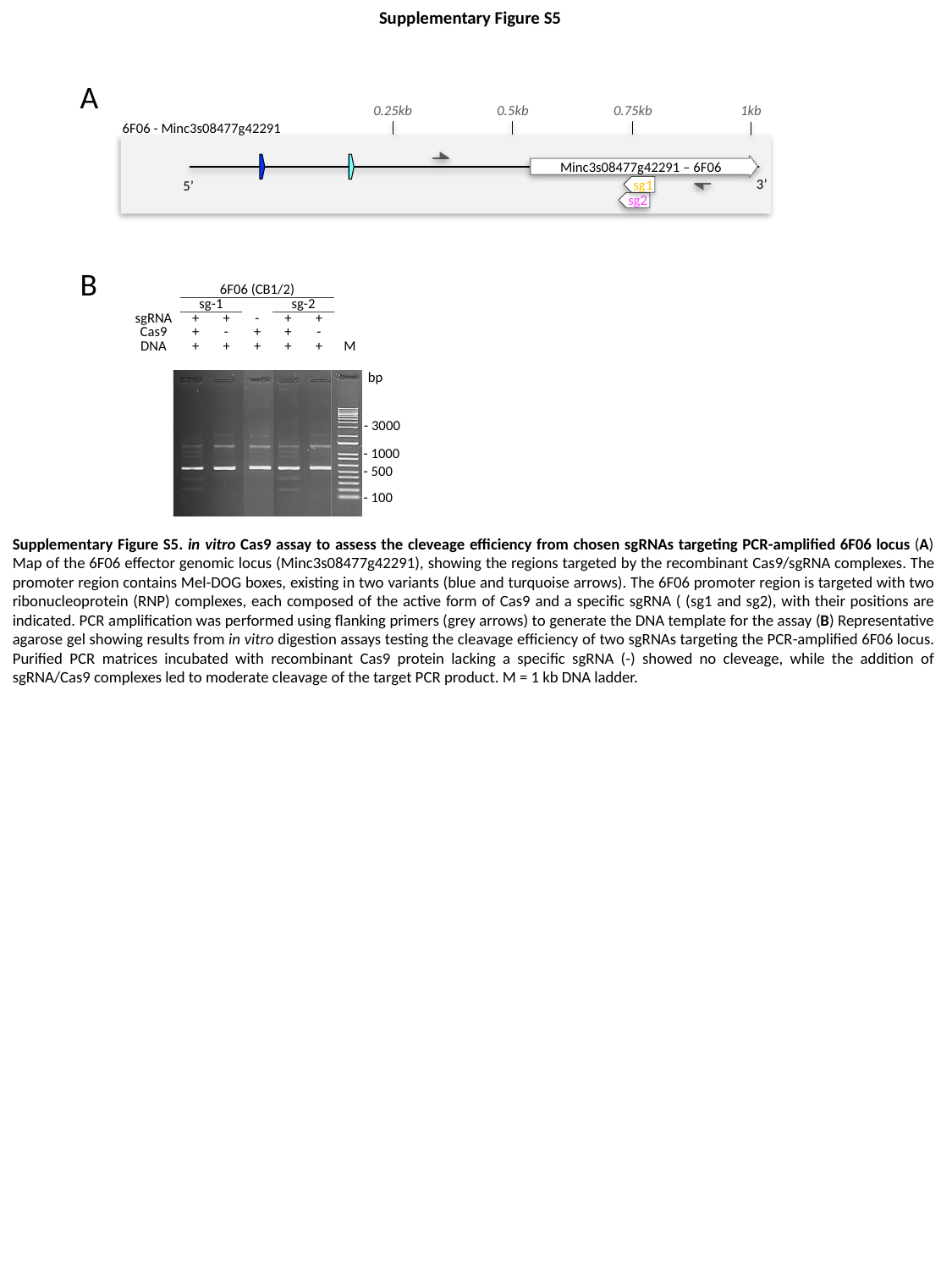

Supplementary Figure S5
A
0.25kb
0.5kb
0.75kb
1kb
6F06 - Minc3s08477g42291
Minc3s08477g42291 – 6F06
3’
sg1
5’
sg2
B
| | 6F06 (CB1/2) | | | | | |
| --- | --- | --- | --- | --- | --- | --- |
| | sg-1 | | | sg-2 | | |
| sgRNA | + | + | - | + | + | |
| Cas9 | + | - | + | + | - | |
| DNA | + | + | + | + | + | M |
bp
- 3000
- 1000
- 500
- 100
Supplementary Figure S5. in vitro Cas9 assay to assess the cleveage efficiency from chosen sgRNAs targeting PCR-amplified 6F06 locus (A) Map of the 6F06 effector genomic locus (Minc3s08477g42291), showing the regions targeted by the recombinant Cas9/sgRNA complexes. The promoter region contains Mel-DOG boxes, existing in two variants (blue and turquoise arrows). The 6F06 promoter region is targeted with two ribonucleoprotein (RNP) complexes, each composed of the active form of Cas9 and a specific sgRNA ( (sg1 and sg2), with their positions are indicated. PCR amplification was performed using flanking primers (grey arrows) to generate the DNA template for the assay (B) Representative agarose gel showing results from in vitro digestion assays testing the cleavage efficiency of two sgRNAs targeting the PCR-amplified 6F06 locus. Purified PCR matrices incubated with recombinant Cas9 protein lacking a specific sgRNA (-) showed no cleveage, while the addition of sgRNA/Cas9 complexes led to moderate cleavage of the target PCR product. M = 1 kb DNA ladder.
